# CysK2 couples copper sensing to metabolic and redox adaptation in *Mycobacterium tuberculosis*

**DOI:** 10.64898/2026.04.16.718964

**Authors:** Wendy Le Mouëllic, Florence Levillain, Ting-Di Wu, Gérald Larrouy-Maummus, Frédéric Lagarrigue, Edith Vidal, Yoann Rombouts, Yannick Poquet, Olivier Neyrolles

**Affiliations:** Institut de Pharmacologie et de Biologie Structurale (IPBS), Université de Toulouse, CNRS, Toulouse, France; Institut Curie, CNRS Unité d’Appui et de Recherche UAR2016, Inserm Unité de Service US43, Université Paris-Saclay, Multimodal Imaging Center, Orsay 91401, France; Centre for Bacterial Resistance Biology, Department of Life Sciences, Faculty of Natural Sciences, Imperial College London, London, United Kingdom

**Keywords:** *Mycobacterium tuberculosis*, copper sensing, cysteine biosynthesis, redox metabolism, intracellular adaptation

## Abstract

Copper is increasingly recognized as a host-derived cue encountered by *Mycobacterium tuberculosis* and other microbes during infections, yet the magnitude, intracellular distribution, and physiological consequences of this exposure remain incompletely understood. Here, we combined high-resolution imaging, transcriptomic profiling, intracellular reporter assays, isotope tracing, and mouse infection models to define how *M. tuberculosis* responds to physiologically relevant copper levels during macrophage infection. NanoSIMS analysis showed that copper reaches intracellular bacilli and accumulates in discrete phosphorus-rich foci in bacteria. Exposure to physiological copper concentrations *in vitro* triggered a highly specific transcriptional response dominated by the copper-inducible CsoR and RicR regulons. The RicR-regulated gene *cysK2* gene, encoding the *S*-sulfocysteine synthase CysK2, was one of the most strongly induced loci. In infected macrophages, *cysK2* expression was modulated by extracellular copper availability, host copper transport pathways, and hypoxia. *In vitro*, an H37Rv *cysK2*-deficient mutant showed reduced amino acid biosynthesis in response to copper exposure. *In vivo*, the mutant was impaired in long-term persistence in mice and displayed a higher oxidation status. Together, these findings identify CysK2 as a copper-responsive metabolic effector that couples host-derived copper sensing to redox homeostasis and intracellular adaptation in *M. tuberculosis*.

**IMPORTANCE:** Successful infection by *Mycobacterium tuberculosis* depends on its ability to detect and adapt to host-imposed changes in the phagosomal environment. Our work shows that copper contributes to this adaptation not simply by imposing toxicity, but by triggering a metabolic response that helps sustain bacterial fitness within host cells. We identify CysK2 as a key component of this response, linking copper sensing to sulfur metabolism, redox balance, and persistence during infection. These findings shift the view of copper from a purely antimicrobial factor to a host-derived environmental signal that remodels *M. tuberculosis* physiology, and they uncover a new mechanism by which the pathogen maintains intracellular survival.

## INTRODUCTION

Tuberculosis (TB), caused by *Mycobacterium tuberculosis*, remains one of the world’s deadliest infectious diseases. In 2024, an estimated 10.7 million new cases and 1.23 million deaths were reported worldwide (1). Despite decades of intensive research and the availability of antimicrobial therapy, *M. tuberculosis* continues to pose a profound global health burden. Its success as a pathogen depends in part on its ability to withstand host-imposed stresses and to remodel its metabolism during infection. Defining these adaptive mechanisms may reveal vulnerabilities that can be exploited to improve TB treatment.

Successful persistence of *M. tuberculosis* within the host requires continuous adaptation to the restrictive intracellular environment, particularly within macrophages. In the phagosome, *M. tuberculosis* encounters multiple stresses, including toxic antimicrobial factors and limitation of essential nutrients (2, 3). Metal ions represent a particularly acute challenge because they are indispensable for bacterial physiology yet become toxic when present in excess. Accordingly, *M. tuberculosis* must tightly control intracellular metal homeostasis, while macrophages exploit metal imbalance as an antimicrobial strategy by withholding essential metals or promoting their toxic accumulation (4–6).

Among metal-dependent host defenses, copper has emerged as a contributor to macrophage antimicrobial activity and is implicated in killing of several bacterial species, including *Escherichia coli* and *Salmonella enterica* serovar Typhimurium (7, 8). In macrophages, copper is imported through CTR1 and can be redistributed by intracellular trafficking pathways involving copper chaperones, such as ATOX1, and the copper-transporting ATPase ATP7A (9–12). Under activating conditions, including lipopolysaccharide (LPS)/interferon (IFN)-γ stimulation, bacterial infection, and hypoxia, *Ctr1* and *Atp7a* expression increases, intracellular copper accumulates while ATP7A relocalizes to vesicular compartments that can associate with pathogen-containing phagosomes. These observations support the idea that macrophages mobilize copper as part of their antimicrobial response (7, 8, 13, 14).

In TB, several observations are consistent with a role for copper during infection. Copper supplementation in drinking water attenuates pulmonary pathology in *M. tuberculosis*-infected mice, although it does not reduce bacterial burden, and copper accumulates within granulomas in the lungs of infected guinea pigs (15). At the cellular level, however, copper dynamics in *M. tuberculosis*-infected macrophages remains poorly defined, as only a single study has reported a transient increase in copper within *M. tuberculosis*-containing phagosomes early after infection (16).

At the bacterial level, the response of *M. tuberculosis* to copper stress has been extensively characterized *in vitro* (17–21). Copper induces a transcriptional program controlled primarily by the copper-responsive repressors CsoR and RicR (22, 23). Notably, the CsoR regulon includes an operon encoding CsoR itself together with CtpV, a putative copper-exporting P-type ATPase, and PacL3, a cognate metal-binding chaperone potentially implicated in resistance to copper intoxication (22–26). While the RicR regulon encodes several proteins involved in copper detoxification, including the metallothionein MymT and the multicopper oxidase MmcO (22, 27, 28), deletion of either or both does not attenuate *M. tuberculosis* growth in mice. However, constitutive repression of all RicR-regulated genes attenuates *M. tuberculosis* fitness *in vivo* (29).

Among the most strongly copper-induced loci in the RicR regulon is the *lpqS-cysK2-rv0849-rv0850* operon, whose function remains poorly understood (22). Of the four predicted gene products encoded by this locus, only an *lpqS* mutant was examined and has no phenotype in copper (29). The downstream-encoded enzyme CysK2 was determined to be an *O*-phosphoserine-dependent cysteine/*S*-sulfocysteine synthase (30), and its contribution to copper adaptation has not been tested.

In this study, we investigated how copper encountered during macrophage infection shapes the adaptation of *M. tuberculosis*. We show that copper reaches intracellular bacilli and induces expression of *cysK2*. Using isotopic labelling, redox reporter strains, and mouse models, we demonstrate that CysK2 sustains amino acid biosynthesis, preserves redox homeostasis, and promotes bacterial persistence during late-stage infection. Collectively, our findings identify CysK2 as a copper-responsive metabolic node linking metal sensing to intracellular adaptation and survival.

## RESULTS

### Copper accumulates in discrete phosphorus-rich foci within intracellular *M. tuberculosis*

Copper accumulation in pathogen-containing vacuoles is widely viewed as part of the macrophage antimicrobial response. However, direct visualization of copper within *M. tuberculosis*-containing compartments is limited. Previous X-ray fluorescence (XRF)-based analysis detected copper in mycobacterial phagosomes, but its spatial resolution did not allow precise subcellular localization within infected cells or bacilli (16).

To determine whether copper reaches intracellular *M. tuberculosis* and to define its subcellular distribution in infected macrophages, we performed high-resolution nanoSIMS imaging. Murine bone marrow-derived macrophages were infected with ^13^C-labeled *M. tuberculosis*, allowing simultaneous detection of copper and ^13^C-enriched bacterial regions. Three days postinfection, we did not detect copper enrichment in the host cell cytosol (Fig. 1, columns 1 and 2). In contrast, copper accumulation was apparent within ^13^C-enriched bacterial regions (Fig. 1, columns 1 and 2). Copper distribution within bacilli was heterogeneous, with most intracellular bacteria containing one or several highly enriched foci. These foci colocalized with phosphorus-rich areas (Fig. 1, columns 3 and 4), consistent with the presence of polyphosphate granules, which have long been described in *M. tuberculosis* (31). Together, these data show that copper reaches intracellular *M. tuberculosis* and accumulates in discrete phosphorus-rich intrabacterial foci.

**Fig 1.**
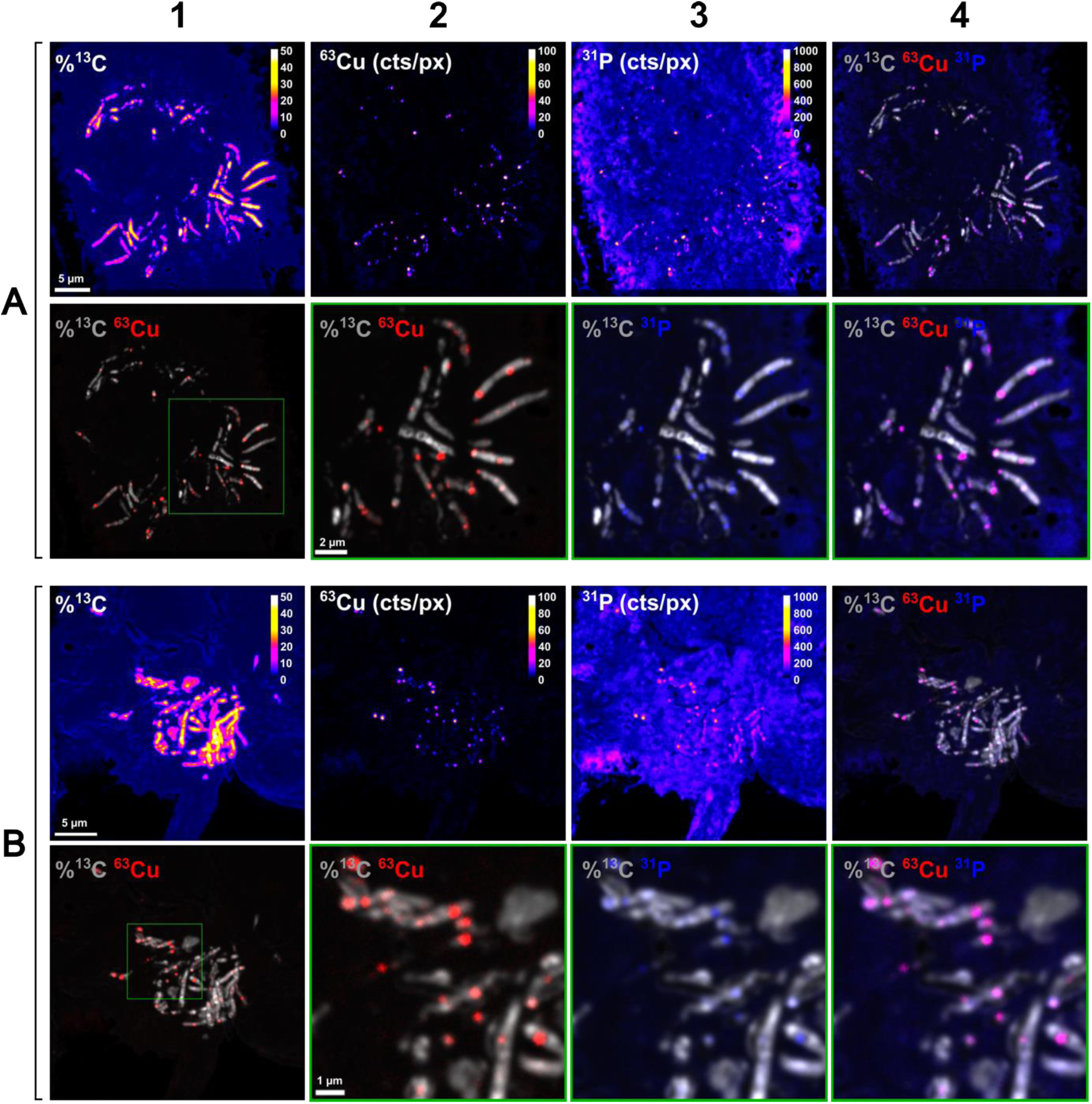
Copper accumulates in discrete phosphorus-rich foci within intracellular *M. tuberculosis*. Murine bone marrow-derived macrophages were infected with *M. tuberculosis* previously cultured in 7H9 supplemented with ^13^C-glycerol. Three days postinfection, cells were fixed with 4% paraformaldehyde and analyzed by nanoSIMS to visualize the distribution of ^13^C, ^63^Cu, and ^31^P in host cells and intracellular bacilli. **(A)** and **(B)** show representative images from two infected macrophages. % ^13^C represent ^13^C atom fraction (%) calculated as ^13^C/(^13^C + ^12^C) × 100. The ^63^Cu and ^31^P signals represent the number of counts per pixel (cts/px).

### Physiological copper levels trigger strong *cysK2* induction in *M. tuberculosis*

Having established that copper can reach intracellular *M. tuberculosis*, we next asked how these physiologic copper levels affected bacterial gene expression. Previous transcriptomic studies examined the response of *M. tuberculosis* to moderate-to-high copper concentrations *in vitro*, but the effects of lower concentrations, such as those reported in phagosomes at 24 h postinfection (16), were unknown. We therefore performed RNA-seq analysis on bacilli exposed to either 20 µM copper sulfate, chosen to approximate physiological exposure (32, 33), or 100 µM (high) copper.

Exposure to 20 µM copper elicited a highly specific transcriptional response (Fig. 2A). Notably, nearly all genes induced at 20 µM copper belong to the CsoR and RicR regulons (Fig. 2B). By contrast, 100 µM copper induced a broad stress response involving more than 50 genes mostly involved in Fe-S cluster biogenesis, mycobactin biosynthesis, and cell envelope-associated functions (Fig. 2B).

**Fig. 2.**
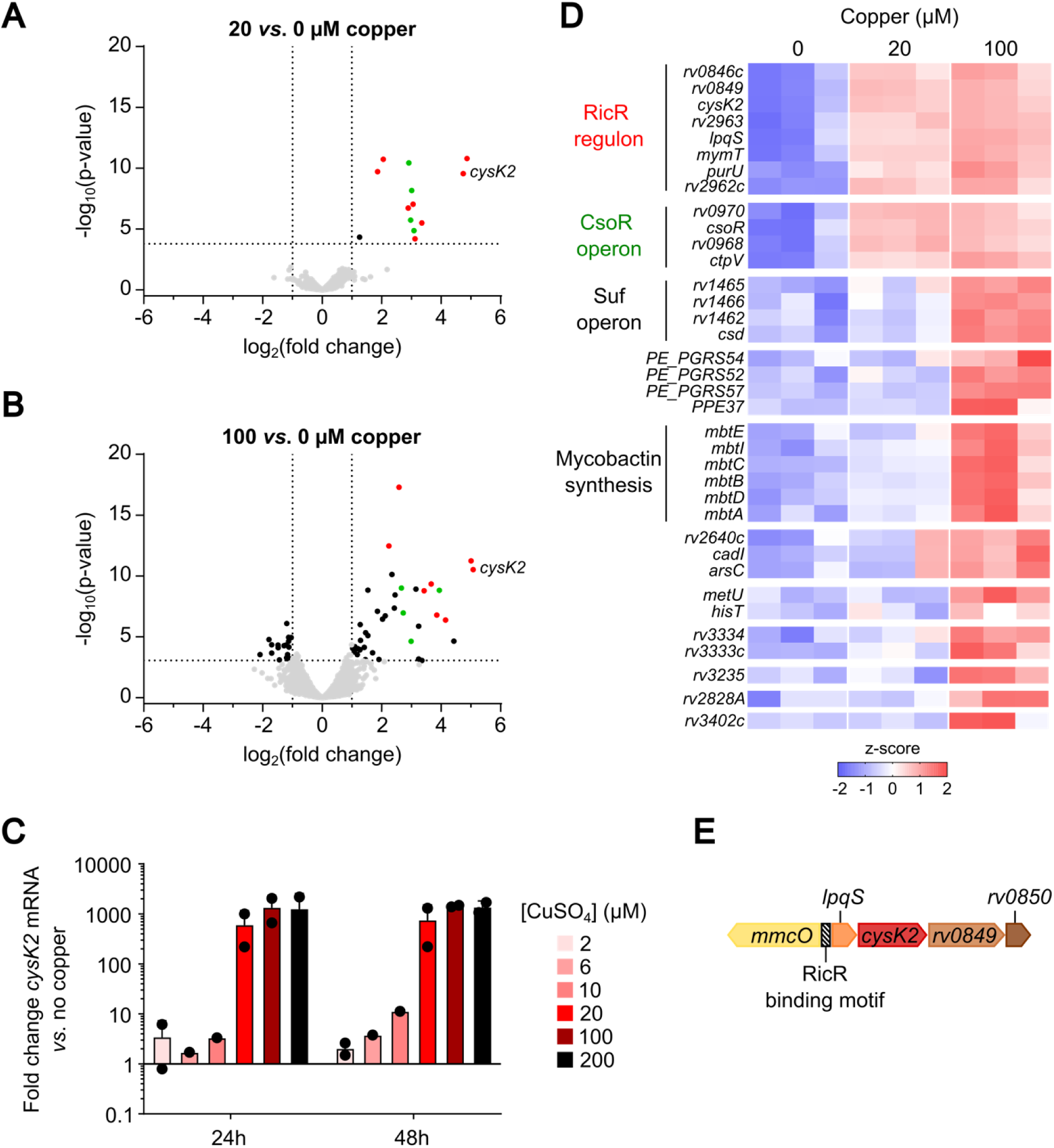
Physiological copper concentrations trigger a focused transcriptional response and strongly induce *cysK2*. Total RNA was isolated from *M. tuberculosis* cultures grown to an OD_600_ of 0.8 in the presence of 0, 20 or 100 µM copper and subjected to RNA-seq analysis. **(A)** Volcano plots showing the distribution of genes according to log_2_(fold change) and - log_2_(P value). **(B)** Heatmaps showing expression changes across selected functional categories. Heatmaps were generated using row z-scores of normalized logCPM values. **(C)** *M. tuberculosis* was incubated in M9 minimal medium supplemented with 0, 2, 6, 10, 20, 100, or 200 µM CuSO_4_. After 24 or 48 h, total RNA was extracted and *cysK2* expression was quantified by RT-qPCR. Data represent mean ± SD fold change relative to the reference gene *rpoB* and the 0 µM CuSO_4_ condition from two independent experiments. **(D)** Genetic organization of the *cysK2* locus in *M. tuberculosis*.

Among the genes most strongly induced at 20 µM copper were *mymT*, *mmcO*, *ctpV*, and *cysK2* (*rv0848*). Because the role of CysK2 in copper adaptation had not been previously investigated, we examined its regulation in more detail. RT-qPCR analysis showed that *cysK2* expression increased approximately 4-fold after exposure to as little as 6 µM copper for 48 h and reached substantial induction, exceeding 500-fold, at 20 µM copper (Fig. 2C). Consistently, a dual reporter strain carrying a constitutive *lpqSp*-mCherry reporter strain confirmed promoter activation at copper concentrations as low as 6 µM (Fig. S1A).

In agreement with a previous report (22), and with the organization of *cysK2* within the RicR-regulated operon (Fig. 2D), other operon-associated genes, including *lpqS*, *rv0849*, and the upstream gene *mmcO*, were also induced by copper (Fig. S1B). Interestingly, the expression of the CsoR-regulated *ctpV* gene also increased in response to copper, albeit its maximal induction was 100-fold lower than the *lpqS-cysK2-rv0849* locus (Fig. S1B). We then asked whether this response extended more broadly to sulfur assimilation and cysteine biosynthesis pathways. However, genes encoding the sulfate transporter SubI (34, 35), the alternative cysteine synthases CysK1 and CysM (36–38), and the reverse transsulfuration enzyme MetB (39) did not show induction comparable to *cysK2* (Fig. S1B). Thus, physiological copper exposure does not broadly induce sulfur metabolic pathways in *M. tuberculosis*, but instead elicits a focused transcriptional response in which CysK2 emerges as a uniquely and strongly induced component of the sulfur metabolic network.

### *cysK2* is genomically associated with copper-homeostasis loci in Actinomycetes

To further explore the potential function of CysK2, we examined its phylogenetic distribution across bacterial phyla using BLAST searches in the UniProt database. Homologs sharing more than 50% amino acid identity with *M. tuberculosis* CysK2 were detected exclusively within Actinomycetes. Within this class, homologs were most frequently found in Kitasatosporales, Mycobacteriales, Pseudonocardiales, and Streptosporangiales, were less common in Micromonosporales, and were rare or absent in other actinomycete lineages (Table S1).

In a representative panel of *Mycobacterium* species spanning rapid- and slow-growing taxa as well as pathogenic, opportunistic, and non-pathogenic species, CysK2 formed a distinct clade separate from the *M. tuberculosis* sulfur-metabolism enzymes CysK1, CysM, Cbs, and Cds1 (Fig. 3A). Within this set, CysK2 was present in species including *M. marinum, M. kansasii, M. fortuitum,* and *M. abscessus*, but absent from others, including *M. avium, M. smegmatis, M. gordonae, M. aurum, M. terrae,* and *M. vaccae* (Fig. 3A). Thus, its presence did not segregate clearly with growth rate or pathogenicity.

**Fig. 3.**
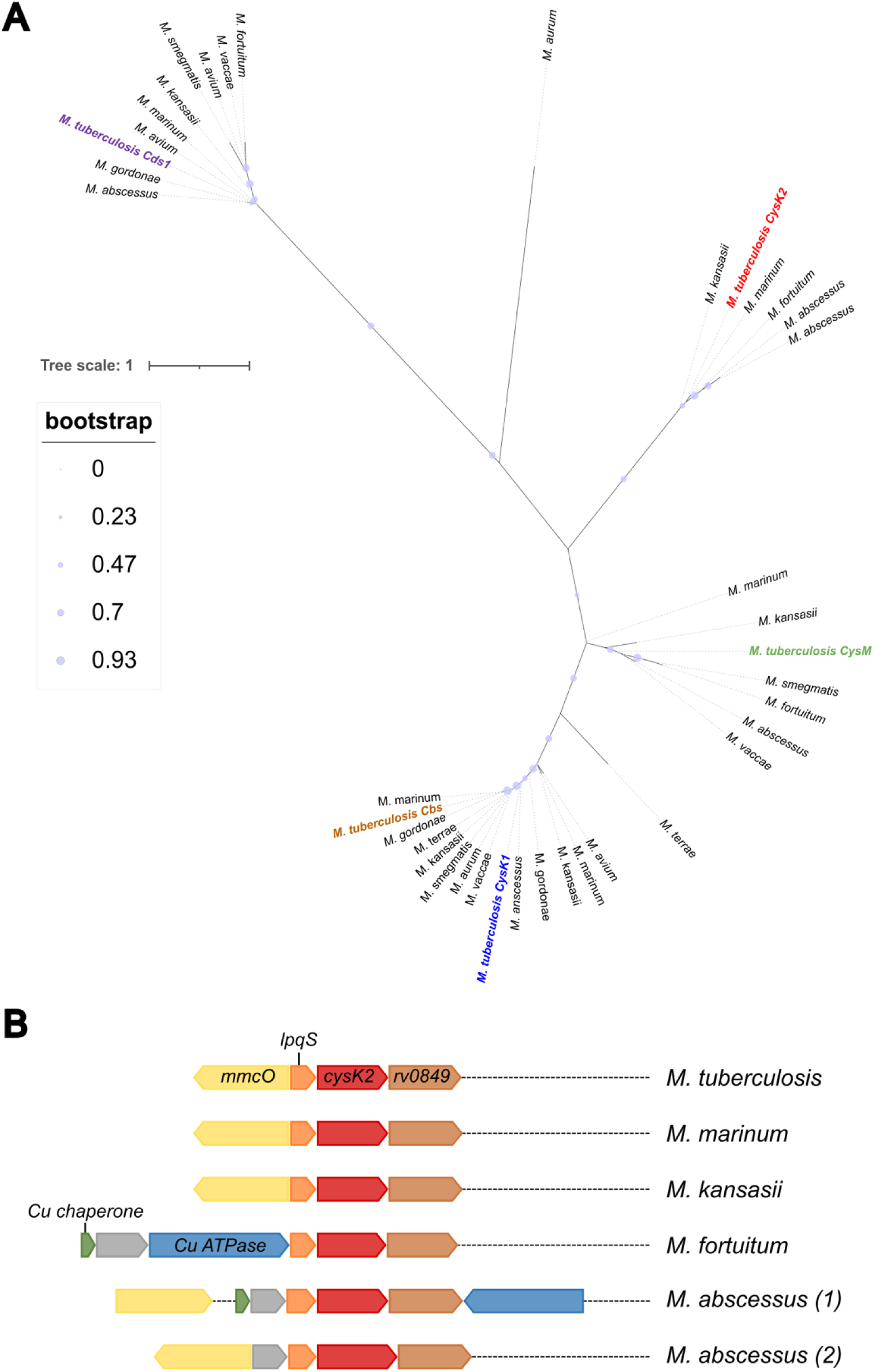
Phylogenetic distribution and genomic context of CysK2-related proteins in Actinomycetes. **(A)** Phylogenetic tree of selected proteins related to the *M. tuberculosis* cysteine synthases CysM, CysK1, Cbs, CysK2, and Cds1. The *M. tuberculosis* reference proteins are highlighted in color: CysM (green), CysK1 (blue), Cbs (maroon), CysK2 (red), and Cds1 (purple). Representative homologs from mycobacteria are shown. The tree was rooted using an *E. coli* homologue. Circle size indicates bootstrap support, as shown in the inset. The scale bar represents 1 substitution per 100 amino acid positions. Protein accession numbers (UniProt) used to construct the tree are listed in Table S2. **(B)** Representative genomic neighbourhoods of *cysK2*-containing loci in selected mycobacteria. Gene arrows indicate predicted coding sequences and transcriptional orientation. Color coding highlights conserved functional categories associated with the locus.

We next examined the genomic context of *cysK2* in the selected CysK2-containing *Mycobacterium* species using the Integrated Microbial Genomes and Microbiomes database. In all species examined, *cysK2* was associated with a conserved copper-related locus (Fig. 3B). Together, these analyses identify *cysK2* as a lineage-restricted actinomycete gene recurrently associated with copper-related genomic neighborhoods. Combined with its strong induction by physiological copper levels in *M. tuberculosis*, this organization supports a role for CysK2 in copper-responsive adaptation.

### Host-derived copper induces *cysK2* expression in intracellular *M. tuberculosis*

We next tested if intracellular copper modulates *cysK2* expression during macrophage infection. Murine macrophages were infected with the *lpqSp*-mCherry reporter strain and increasing concentrations of copper were added to culture media after bacterial internalization. Four days postinfection, *lpqSp* activity was quantified as the mCherry/GFP fluorescence ratio in intracellular bacilli. No significant increase in reporter activity was detected below 20 µM extracellular copper, whereas higher concentrations triggered a progressive increase in *lpqSp* activity that did not saturate even at 200 µM (Fig. 4A). In contrast to the strong and saturable induction observed in broth culture at concentrations as low as 6 µM copper (Fig. S1A), these data indicate that only a fraction of extracellular copper reaches intracellular bacilli during macrophage infection.

**Fig. 4.**
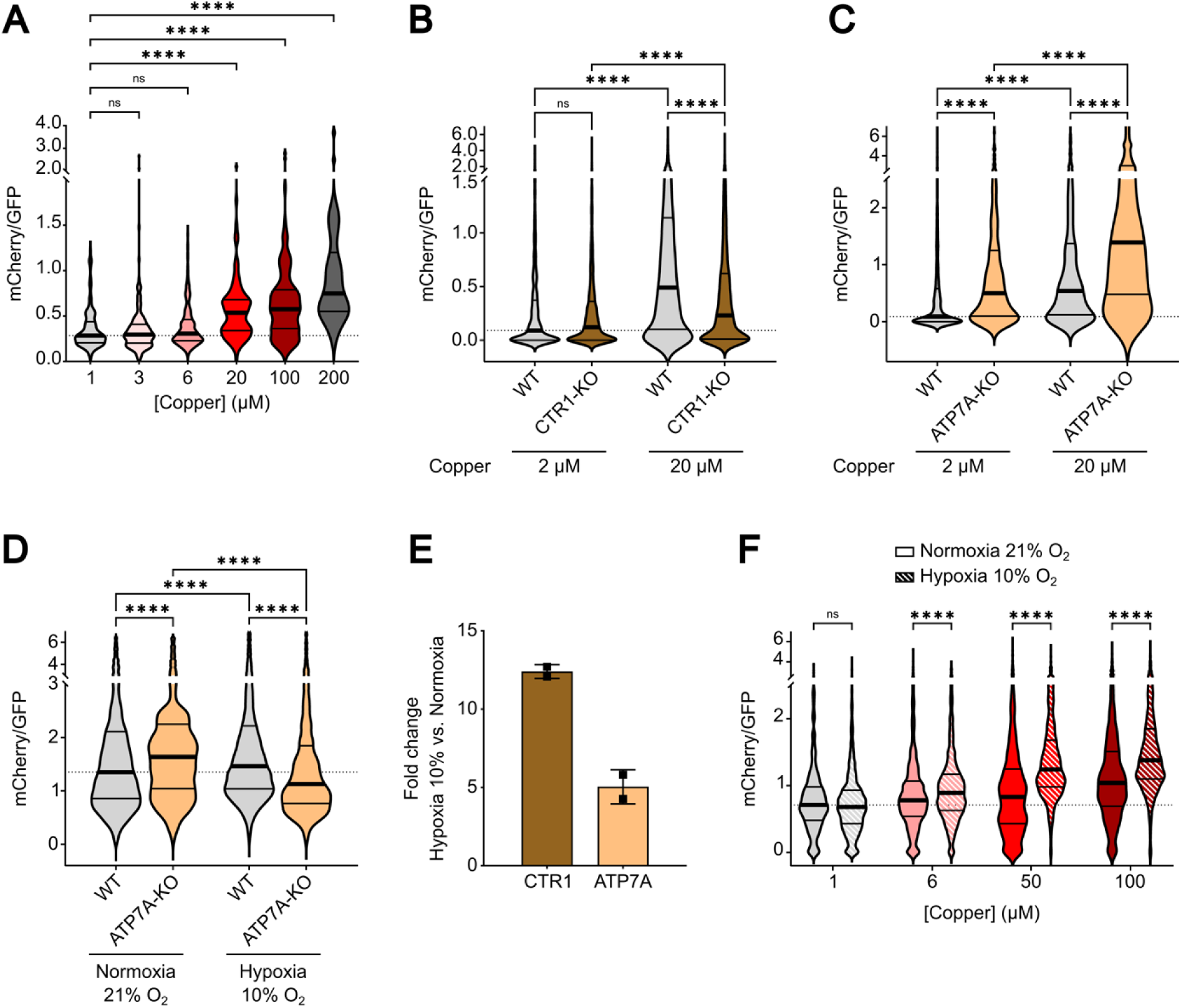
Host-derived copper induces *cysK2* expression in intracellular *M. tuberculosis*. **(A)** Murine bone marrow-derived macrophages (BMDMs) were infected with the *M. tuberculosis* P1-GFP *lpqSp-*mCherry reporter strain in the presence of increasing extracellular copper concentrations (CuCl_2_). At 4 days postinfection, cells were fixed and *lpqSp* activity in intracellular bacilli was quantified by confocal microscopy as the mCherry/GFP fluorescence ratio. Violin plots show individual bacteria (n = 38-264) from three wells and are representative of two independent experiments. Statistical analysis: Kruskal-Wallis test followed by Dunn’s multiple-comparison test. **(B-C)** Wild-type, CTR1-knockout, or ATP7A-knockout HoxB8-derived macrophages were infected with the reporter strain under basal copper conditions (2 µM, contributed by 10% FBS) or after supplementation with 20 µM CuSO_4_. At 6 days postinfection, *lpqSp* activity in intracellular bacilli was quantified as the mCherry/GFP fluorescence ratio. Violin plots show individual bacteria (n = 739-3,735) from two or three wells and are representative of two independent experiments. Statistical analysis: Kruskal-Wallis test followed by Dunn’s multiple-comparison test. **(D)** Wild-type and ATP7A-knockout HoxB8-derived macrophages were infected with the reporter strain and cultured with 20 µM CuSO_4_ under normoxic (21% O_2_) or hypoxic (10% O_2_) conditions. At 6 days postinfection, *lpqSp* activity in intracellular bacilli was quantified as the mCherry/GFP fluorescence ratio. Violin plots show individual bacteria. Statistical analysis: Kruskal-Wallis test followed by Dunn’s multiple-comparison test. **(E)** BMDMs were activated with IFN-γ and LPS and cultured under normoxic (21% O_2_) or hypoxic (10% O_2_) conditions for 3 days. Total RNA was extracted, and *Ctr1* and *Atp7a* expression was quantified by RT-qPCR. Data represent mean ± SD from two independent RNA extractions. **(F)** Murine BMDMs were infected with the reporter strain in the presence of increasing extracellular copper concentrations (CuSO_4_) under normoxic (21% O_2_) or hypoxic (10% O_2_) conditions. At 4 days postinfection, *lpqSp* activity in intracellular bacilli was quantified by confocal microscopy as the mCherry/GFP fluorescence ratio. Violin plots show individual bacteria (n = 89-1,892) from two wells and are representative of two independent experiments. Statistical analysis: Kruskal-Wallis test followed by Dunn’s multiple-comparison test.

To determine whether this response depends on host copper trafficking, we infected wild-type, CTR1-, or ATP7A-KO HoxB8-derived macrophages with the reporter strain and measured *lpqSp* activity under basal (2 µM) or physiological (20 µM) copper conditions (Fig. S2). Six days postinfection, copper supplementation increased reporter activity in all three cell lines (Fig. 4B, 4C), consistent with the results obtained in primary macrophages. However, *lpqSp* activity was reduced in intracellular bacilli residing in CTR1-deficient macrophages compared with wild-type cells (Fig. 4B), consistent with the role of CTR1 in extracellular copper import. By contrast, bacilli infecting ATP7A-deficient macrophages displayed higher reporter activity than those in wild-type cells under the same conditions (Fig. 4C), indicating that ATP7A influences copper availability to intracellular bacteria in a reverse manner. Together, these results show that host-derived copper can induce *cysK2* expression in intracellular *M. tuberculosis* and that this response is modulated by macrophage copper transport pathways.

### Hypoxia enhances host-dependent copper signalling tointracellular *M. tuberculosis*

Because ATP7A relocalizes in response to conditions associated with macrophage activation or hypoxia, a feature of the deep lung and of TB lesions, we next examined its contribution under reduced oxygen tension. We infected wild-type and ATP7A-deficient macrophages with the *lpqSp* reporter strain and cultured at 10% O_2_, a level comparable to that reported in lung tissue. Under these conditions, reporter activity was lower in bacilli residing in ATP7A-deficient macrophages than in wild-type cells (Fig. 4D), indicating that ATP7A was required for maximal copper-dependent *cysK2* induction under hypoxia.

Consistent with this result, 10% O_2_ increased *Ctr1* and *Atp7a* expression in LPS/IFN-γ-activated primary macrophages relative to normoxia (Fig. 4E), and infected macrophages cultured at 10% O_2_ showed increased *lpqSp* activity compared with normoxic controls at the same extracellular copper concentration (Fig. 4F). By contrast, planktonic bacilli exposed to 10% O_2_ did not show increased reporter activity across copper concentrations (Fig. S3), indicating that the effect of hypoxia is mediated by the host rather than by a direct bacterial response to this oxygen level. These results indicate that moderate tissue-like hypoxia enhances copper-dependent *cysK2* induction in intracellular *M. tuberculosis* through a host-dependent mechanism involving ATP7A and CTR1.

### CysK2 promotes redox homeostasis and persistence of *M. tuberculosis in vivo*

To determine whether CysK2 contributes to *M. tuberculosis* virulence, we first assessed bacterial fitness during infection of susceptible C3HeB/FeJ mice, which develop granuloma-like structures that do not form in C57BL/6 mice (40). We infected animals by aerosol with a low-to-medium bacterial dose (∼100 CFU), and monitored pulmonary bacterial burdens over time (Fig. 5A). The Δ*cysK2* mutant (Fig. S4) showed moderate but significant attenuation during late-stage infection compared with wild-type and complemented strains, indicating that CysK2 contributes to bacterial persistence *in vivo*. To determine whether this phenotype was host-model specific, we performed parallel infections in resistant C57BL/6 mice. Consistent with the findings in C3HeB/FeJ animals, the Δ*cysK2* strain displayed a reduction in lung bacterial loads at late time points (Fig. S5), confirming a reproducible role for CysK2 in long-term infection.

**Fig. 5.**
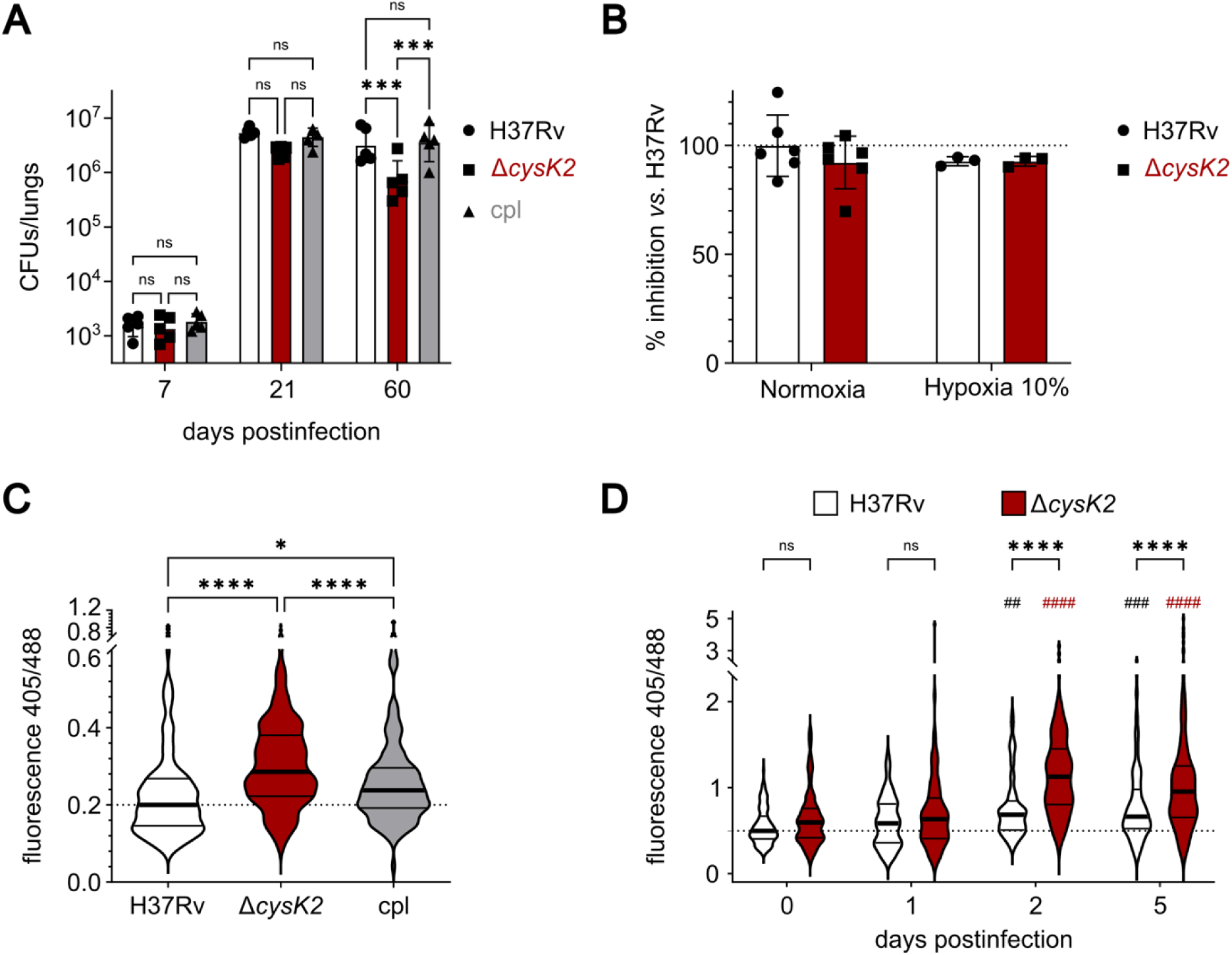
CysK2 promotes *M. tuberculosis* persistence and redox homeostasis during infection. **(A)** C3HeB/FeJ mice were infected by aerosol with H37Rv, Δ*cysK2*, or the complemented strain (∼100 CFU per mouse). Lungs were collected at 7, 21, and 60 days postinfection, and bacterial burden was quantified by CFU enumeration. Data represent mean ± SD for n = 5 mice per strain. Statistical analysis: two-way ANOVA with Tukey’s post hoc test on log-transformed data. **(B)** Copper sensitivity of H37Rv and Δ*cysK2* strains was assessed by disk diffusion assay using 5 µL of 1 M CuSO_4_. Plates were incubated under normoxic (21% O_2_) or hypoxic (10% O_2_) conditions, and inhibition zones were measured and normalized to the H37Rv wild-type strain under normoxia. Data represent mean ± SD of three or six biological replicates from two independent experiments. **(C)** C3HeB/FeJ mice were infected with H37Rv, Δ*cysK2*, and complemented strains expressing the Mrx1-roGFP redox reporter. At 60 days postinfection, lungs were collected and analyzed by confocal microscopy. The 405/488 fluorescence ratio was quantified in individual bacteria as a measure of redox state. Violin plots show median and interquartile range for n = 115 (H37Rv), n = 341 (Δ*cysK2*), and n = 124 (complemented) bacteria. Statistical analysis: Kruskal-Wallis test followed by Dunn’s multiple-comparison test. **(D)** Murine bone marrow-derived macrophages were infected with H37Rv or Δ*cysK2* strains expressing the Mrx1-roGFP reporter. At the indicated times postinfection, cells were fixed and the 405/488 fluorescence ratio of intracellular bacteria was quantified by confocal microscopy. Violin plots show median and interquartile range for n = 88-131 individual bacteria. Statistical analysis: two-way ANOVA with Tukey’s post hoc test. ## indicates comparison with day 0 for the corresponding strain

Because *cysK2* is strongly induced by copper during macrophage infection, particularly under tissue-like hypoxia, we next asked whether CysK2 contributes to copper resistance. We therefore compared the copper sensitivity of wild-type and Δ*cysK2* strains *in vitro* using disk diffusion assays (Fig. 5B), growth measurements in copper-supplemented rich medium (Fig. S6A), and CFU enumeration after copper exposure (Fig. S6B). Given that copper toxicity can be enhanced under low-oxygen conditions (41–44), we also tested sensitivity at 10% O_2_ (Fig. 5B). Under all conditions examined, the H37Rv Δ*cysK2* mutant behaved similarly to the wild type, suggesting that the *in vivo* attenuation of the Δ*cysK2* strain was not explained by increased copper sensitivity, at least in a H37Rv background.

We next investigated whether CysK2 contributes to redox homeostasis during infection. To assess bacterial redox status *in vivo*, we generated wild-type, Δ*cysK2*, and complemented strains expressing the Mrx1-roGFP reporter, which monitors intracellular redox changes through shifts in the 405/488-nm excitation ratio (45). We infected C3HeB/FeJ mice with these strains, and analyzed lung sections by confocal microscopy 60 days postinfection (Fig. 5C). The oxidized/reduced fluorescence ratio was significantly higher in Δc*ysK2* bacilli than in wild-type and complemented strains, indicating increased oxidative stress during chronic infection. A similar redox imbalance was observed *ex vivo* in infected macrophages, where Δ*cysK2* bacteria exhibited a more oxidized state as early as 2 days postinfection (Fig. 5D). Together, these data show that CysK2 promotes *M. tuberculosis* persistence during late-stage infection and supports redox homeostasis *in vivo* and in infected macrophages.

### CysK2 supports amino acid biosynthesis during copper exposure

Because CysK2 is predicted to contribute to cysteine production, we tried to assess whether intracellular cysteine levels differed between wild type H37Rv and Δ*cysK2* bacilli. Cytosolic cysteine was analyzed by LC/MS and GC/MS under conditions that induce *cysK2* expression. However, cysteine remained undetectable in both strains, consistent with previous reports indicating that intracellular cysteine levels in *M. tuberculosis* are extremely low or below the detection limit (46).

We next assessed the impact of CysK2 on amino acid biosynthesis using short-term isotope tracing. Wild-type and Δ*cysK2* bacilli were incubated for 4 h with 1 mM ^15^N-serine in the presence of increasing copper concentrations. As expected, the highest ^15^N incorporation was detected in glutamine, which serves as a major nitrogen donor for subsequent transamination reactions (Fig. 6, first panel). In addition to serine, 13 amino acids showed significant ^15^N labelling (Fig. 6). In the wild-type strain, labelling was highest at 6 µM copper and decreased significantly at 20 µM copper. Increasing copper to 100 µM produced only limited additional effects for several amino acids, including Thr, Ile, Phe, Val, Leu, and Glu. By contrast, the Δ*cysK2* mutant incorporated substantially less ^15^N into amino acids than the wild-type strain at both 6 and 20 µM copper. At 100 µM copper, labelling in the mutant remained slightly lower than in wild-type for most amino acids, although the difference was no longer apparent for Tyr, Thr, His, Leu, and Arg. These results show that copper inhibits amino acid biosynthesis in *M. tuberculosis* and that CysK2 helps preserve this process at low and physiologically relevant copper concentrations, as indicated by the reduced ^15^N incorporation observed in the Δ*cysK2* mutant relative to the wildtype. At higher copper concentrations, however, labelling decreased in both strains, suggesting that copper-mediated inhibition eventually exceeds the contribution of CysK2.

**Fig. 6.**
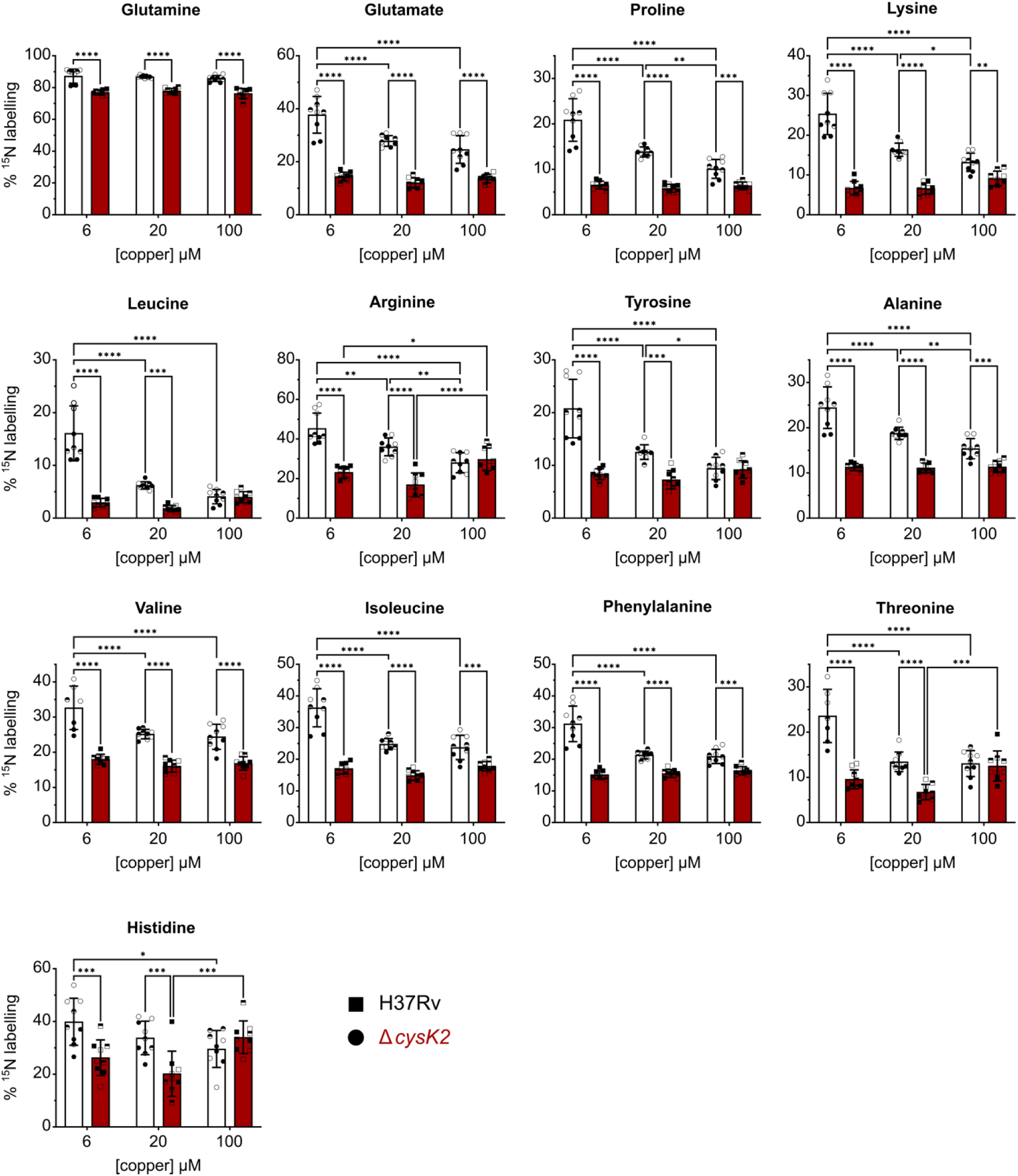
CysK2 supports amino acid biosynthesis during copper exposure. Wild-type *M. tuberculosis* H37Rv and Δ*cysK2* strains were incubated on minimal agar medium supplemented with 1 mM ^15^N-serine for 4 h in the presence of 6, 20, or 100 µM copper. Metabolites were then extracted, and ^15^N incorporation into individual amino acids was quantified by LC-MS. Data represent the mean of three independent cultures, each analyzed in triplicate. Statistical analysis: two-way ANOVA with Tukey’s post hoc test.

## DISCUSSION

Our study shows that host-derived copper reaches intracellular *M. tuberculosis* during macrophage infection and elicits a focused adaptive response rather than a broad stress program. We identify *cysK2* as a major copper-responsive gene whose expression depends on copper availability, host copper trafficking, and hypoxia, and show that CysK2 contributes to redox homeostasis, maintenance of amino acid biosynthetic flux, and long-term persistence *in vivo*. Together, our findings support a model in which copper acts as a host-derived cue that promotes CysK2-mediated metabolic adaptation in intracellular *M. tuberculosis*.

We first sought to follow the path of copper from the extracellular environment to intracellular bacilli. Using high-resolution nanoSIMS, we show that copper reached *M. tuberculosis* within infected macrophages and accumulated in discrete phosphorus-rich intrabacterial foci, most likely corresponding to polyphosphate granules. To our knowledge, this is the first direct evidence in *M. tuberculosis* that copper can concentrate in structures consistent with polyphosphate-dependent sequestration. Copper binding by polyphosphates has previously been described in other systems, including bacteria and the unicellular alga *Chlamydomonas* (47, 48). By contrast, previous scanning transmission electron microscopy (STEM)-based elemental analysis of polyphosphate granules in planktonic mycobacteria identified potassium, calcium, and magnesium, but not copper (49). This discrepancy may reflect the different biological contexts examined, since our analysis was performed on intracellular bacilli, but it may also arise from technical differences in fixation and sample processing that affect retention of labile metal ions.

Our data also refine the current view of copper accumulation in the mycobacterial phagosome. Apart from these intrabacterial phosphorus-rich foci, we did not detect prominent copper enrichment in the surrounding compartment, in line with the earlier XRF study by Wagner and colleagues showing that phagosomal copper levels are modest at later time points after infection, reaching approximately 25 µM at 24 h (16). Importantly, the much higher copper concentration often cited in the literature corresponds to an early and highly variable transient peak measured 1 h after infection (426 ± 393 µM) (16). These observations suggest that intracellular *M. tuberculosis* is not necessarily exposed to sustained, massively elevated copper concentrations throughout infection. Rather, bacilli may experience lower and dynamic copper levels over time, with part of the incoming metal becoming concentrated in discrete intrabacterial structures.

Our RNAseq data are consistent with physiological copper acting as a signal that engages the copper-responsive regulators, CsoR and RicR, rather than inducing a broad intoxication program. Among the induced genes, *cysK2* emerged as one of the most strongly responsive. In the companion paper to this work, Ji *et al*. showed that CysK2 is in fact a cysteine synthase and had been previously mischaracterized as an *O*-phosphoserine-dependent cysteine/*S*-sulfocysteine synthase (30). Crucially, this activity is essential for robust copper resistance in CDC1551 and Erdman *M. tuberculosis* strains (50). In this regard, our phylogenetic and synteny analyses are informative: close CysK2 homologs are restricted to Actinomycetes, and in several lineages *cysK2* is embedded in conserved loci containing copper-homeostasis genes. This lineage-restricted genomic organization argues against a purely housekeeping role and instead strongly supports the idea that CysK2 participates in a specialized adaptive program linked to copper-responsive physiology.

Using a copper-responsive reporter, we further show that *cysK2* expression is modulated by host-derived copper during macrophage infection. Increasing extracellular copper after bacterial internalization resulted in progressively higher *lpqSp* activity in intracellular bacilli. In contrast to the rapid, saturable induction observed in broth culture, this response was not saturated even at the highest extracellular copper concentrations tested, consistent with the idea that only a fraction of extracellular copper is ultimately delivered to intracellular *M. tuberculosis*. These observations suggest that *cysK2* expression may act as a sensitive readout of copper availability in infected cells and, more broadly, that copper exposure of intracellular bacilli depends on the host environment rather than reflecting a fixed, uniformly high phagosomal copper concentration. This point may be important when considering copper biology across infection models. Tissue copper concentrations increase during infection, especially in the lung (26, 51), but the magnitude and distribution of this increase may differ across species and lesion types. For example, copper accumulation has been reported in granulomas of guinea pigs during TB infection, whereas evidence for comparable lesion-level accumulation in mice is less clear (15, 52). Such differences could help explain why copper-responsive genes that contribute to copper resistance *in vitro* or appear relevant in one animal model do not always show the same contribution in another.

Our experiments using copper-transporter-deficient macrophages begin to address how copper reaches intracellular bacilli. The reduced *cysK2* induction observed in CTR1-deficient macrophages is fully consistent with the established role of CTR1 in copper import at the plasma membrane and indicates that host copper uptake contributes directly to the signal sensed by intracellular *M. tuberculosis*. The role of ATP7A appears more complex. Under basal conditions, ATP7A deficiency increased reporter activity, suggesting that ATP7A can limit copper availability to bacilli in this context, perhaps by redistributing copper to compartments other than the phagosome or by influencing whole-cell copper homeostasis. This interpretation is compatible with previous work reporting altered intracellular copper levels in ATP7A-deficient macrophages (53). However, under moderate hypoxia, ATP7A became necessary for maximal *cysK2* induction, consistent with prior reports that hypoxia and macrophage activation increase *Ctr1* and *Atp7a* expression and promote ATP7A relocalization away from the Golgi (8, 14). Thus, copper trafficking to the mycobacterial compartment is not static but depends on the physiological state of the host cell. They also support the idea that hypoxia, a feature of infected lung tissue and granulomatous lesions, can enhance copper mobilization toward intracellular bacilli.

Despite its strong induction by copper, the H37Rv Δ*cysK2* mutant did not display overt copper hypersensitivity in the *in vitro* conditions we tested. This is in line with the companion study by Ji *et al*. (50), which shows that H37Rv CysK2 is a hypomorphic allele carrying a nonconserved Ser_93_Gly substitution. Ji *et al.* further found that CysK2 acts as a cysteine synthase and that the H37Rv G_93_ variant produces cysteine far less efficiently than the ancestral S_93_ enzyme. In agreement with this reduced activity, CysK2_G93_ confers only a subtle copper-resistance phenotype in H37Rv, whereas CysK2_S93_ mediates robust copper resistance in H37Rv, CDC1551, and Erdman. Together, these findings indicate that CysK2-dependent cysteine synthesis contributes to copper adaptation, and suggest that even the weak activity of H37Rv CysK2_G93_ is sufficient to provide a fitness advantage during persistent infection by supporting metabolic and redox homeostasis.

In two mouse models, loss of *cysK2* impaired long-term bacterial persistence, and this defect was accompanied by a more oxidized intracellular state both *in vivo* and in infected macrophages. These findings strongly support a role for CysK2 in redox homeostasis during infection. In line with our results, a recent report likewise linked CysK2 to *M. tuberculosis* survival in macrophages and in mice, although the magnitude of the *in vivo* phenotype differed substantially from that observed here (54). This redox-centered view of CysK2 function is consistent with the known importance of sulfate uptake and sulfur assimilation for *M. tuberculosis* virulence (34, 55) and supports the view that the multiple cysteine-biosynthetic pathways of *M. tuberculosis* make distinct contributions under specific host conditions. In this framework, CysK2 appears to be part of a copper-responsive metabolic adaptation that helps preserve thiol-dependent redox balance during chronic infection.

Our isotope-tracing experiments further clarify how CysK2 may support persistence and redox homeostasis during infection. Direct quantification of intracellular cysteine was not technically possible, consistent with previous reports that cysteine levels in *M. tuberculosis* are extremely low or below the detection threshold (46). However, short-term ^15^N-serine labelling revealed that copper exposure slows amino acid biosynthesis and that this effect is exacerbated in the absence of CysK2, particularly at low and physiologically relevant copper concentrations. Thus, CysK2 appears to help preserve amino acid biosynthesis during copper exposure. One plausible interpretation is that copper-induced expression of *cysK2* helps maintain cysteine production under these conditions, thereby supporting broader biosynthetic activity.

Together, our results support a model in which host-derived copper acts as a cue sensed by intracellular *M. tuberculosis*, inducing CysK2-dependent metabolic and redox adaptation that promotes persistence during infection.

## MATERIALS AND METHODS

### Bacterial strains and culture conditions

*M. tuberculosis* H37Rv, mutant strains, and recombinant derivatives were grown at 37°C in liquid Middlebrook 7H9 medium (Difco) supplemented with 10% albumin-dextrose-catalase (ADC; Difco) and 0.05% Tyloxapol (Sigma), or on solid Middlebrook 7H11 medium (Difco) supplemented with 10% oleic acid-albumin-dextrose-catalase (OADC; Difco). When required, hygromycin (50 µg/mL), streptomycin (25 µg/mL), kanamycin (20 µg/mL), or zeocin (25 µg/mL) was added to the culture medium.

### Construction of *M. tuberculosis* mutant and complemented strain

The Δc*ysK2* mutant was generated in *M. tuberculosis* H37Rv by allelic exchange using recombineering, as previously described (56), with the primers listed in Table S2. The allelic exchange substrate (AES) was assembled by three-fragment PCR and consisted of the upstream region of *cysK2*, a zeocin-resistance cassette flanked by two identical dif sites in the same orientation, and the downstream region of *cysK2* (Fig. S4).

For recombineering, the recipient strain carrying the pJV53H plasmid was grown in 7H9 medium supplemented with 10% ADC, 0.05% Tyloxapol, and hygromycin to exponential phase. Recombineering functions were induced by addition of 0.2% acetamide (Sigma). Electrocompetent bacteria were prepared and electroporated with 100 ng of AES, then allowed to recover for 48 h in antibiotic-free 7H9 medium. Bacteria were plated on 7H11 agar supplemented with zeocin (25 µg/mL) and hygromycin (20 µg/mL). Zeocin-resistant colonies were isolated and grown in liquid medium to allow spontaneous recombination between the two *dif* sites and excision of the zeocin-resistance cassette, which was verified phenotypically (Fig. S4).

The complementation plasmid pGMC-S-*lpqSp*-*lpqS*-*cysK2* was constructed by In-Fusion cloning (Takara) using pDE43-MCS as the destination vector and the primers listed in Table S2. The resulting plasmid was electroporated into the *M. tuberculosis* H37Rv Δ*cysK2*::dif4 strain carrying pJV53H.

Deletion of *cysK2* and complementation were verified by PCR on genomic DNA and by RT-qPCR on reverse-transcribed total RNA using specific primers (Table S2). For deletion, primers flanking the upstream and downstream regions of the locus were used; for complementation, primers internal to *cysK2* were used (Fig. S4).

### Construction of *M. tuberculosis* reporter strains

To generate the *cysK2* promoter reporter plasmid, a fragment of approximately 700 bp upstream of *lpqS* was amplified by PCR using the primers listed in Table S2 and cloned by In-Fusion cloning (Takara) upstream of the mCherry coding sequence in a recipient pGMCS plasmid carrying *gfp* under the control of the constitutive P1 promoter.

To generate the redox reporter construct, the Mrx1 (Rv3198A) coding sequence was amplified from *M. tuberculosis* H37Rv genomic DNA, and the linker-roGFP2 open reading frame was amplified from the plasmid pLPCX cyto Grx1-roGFP2. The resulting PCR fragments were assembled by Gibson assembly into linearized pGEM-5Z. The Mrx1-roGFP2 cassette was then subcloned into the pVV16 vector. Reporter plasmids were electroporated into the indicated *M. tuberculosis* strains, and fluorescence of recombinant clones was verified by flow cytometry.

### Generation and differentiation of HoxB8-immortalized myeloid progenitor cells

Immortalized HoxB8 progenitor cells derived from Cas9-expressing mice were maintained in myeloid medium consisting of RPMI GlutaMAX supplemented with 10% fetal bovine serum (FBS; Merck), antibiotic-antimycotic (Gibco), 20 ng/mL GM-CSF (Miltenyi Biotec), and 0.5 µM β-estradiol. Cells were passaged every 3 days. For macrophage differentiation, HoxB8 progenitors were washed twice with PBS and seeded at 5 × 10^5^ cells per well in 6-well plates in complete medium (RPMI supplemented with FBS and antibiotic-antimycotic) containing 20 ng/mL mouse M-CSF (Miltenyi Biotec). Fresh medium was added on days 3 and 6. Differentiated HoxB8-derived macrophages were harvested by incubation in PBS containing 5 mM EDTA for 5 min at 37°C. For gene targeting, sgRNAs were cloned individually into the pLenti-sgRNA vector (Addgene #71409). An sgRNA targeting luciferase was used as a control. Plasmids encoding the corresponding sgRNAs were prepared using Lipofectamine 3000 (Thermo Fisher) and Opti-MEM (Gibco) according to the manufacturer’s instructions. For lentiviral transduction, 2 × 10^5^ HoxB8 myeloid progenitor cells were incubated in 1 mL myeloid medium containing pLenti-sgRNA lentiviral particles in the presence of LentiBlast Premium (OZ Biosciences). After 24 h, cells were centrifuged at 300 × *g* for 5 min, and the medium was replaced with myeloid medium containing puromycin (7 µg/mL; InvivoGen) for selection for 72 h.

### Proteomics

HoxB8-derived macrophages were cultured as described above in 24 well plates. Cells were lysed in a solution of 5% sodium dodecyl sulfate (SDS) (Invitrogen) and 50 mM triethylammonium bicarbonate, pH 8.5. Protein disulfide bridges and cysteine residues were reduced and alkylated by incubation with 10mMTris (2-carboxyethyl) phosphine hydrochloride (NeoBiotech) and 40mM 2-chloroacetamide (Aldrich), 5min at pH 8.5 and 95°C. Proteins were digested on S-trap Micro devices (Protifi) using the manufacturer’s protocol. Briefly, samples were acidified by addition of phosphoric acid (2.5% final concentration), and proteins were precipitated by addition of 6 volumes of 90% methanol in 100mM TEAB, pH7.5. Samples were applied on the S-Trap Micro devices and precipitated proteins were washed 6 times with 90% methanol, 100mM TEAB, pH7.5. Proteins were then digested by incubation with trypsin (Promega) at 1% (trypsin/protein) and 37°C overnight. The resulting peptides were successively eluted with 40µl of 50mM ammonium bicarbonate, 40µl of 0.2% formic acid and 40µl of 50% acetonitrile, 0.2% formic acid. and speed-vacuum dried. After resuspension in 0.1% formic acid, they were desalted on Affinisep C18 T1 Tips with 0.1% formic acid, eluted with 40% acetonitrile, 0.1% formic acid and then with 70% acetonitrile, 0.1% formic acid, and finally dried again.

Peptides were analyzed by nanoscale liquid chromatography using an UltiMate 3000 system (Thermo Scientific) coupled to an Orbitrap Exploris 480 mass spectrometer (Thermo Scientific). One µl of each sample was injected on the analytical C18 column (75 μm inner diameter × 50 cm, Pepmap C18, 2 µm, Thermofisher Electron SAS) equilibrated in 95% solvent A (5% acetonitrile, 0.2% formic acid) and 5% solvent B (80% acetonitrile, 0.2% formic acid). Peptides were eluted using a 2.5%-40% gradient of solvent B over 62 min at a flow rate of 300 nl/min. The mass spectrometer was operated in data-independent acquisition (DIA) mode with the Xcalibur software. Full scan MS1 scans were acquired on the m/z mass range 400–1008, with a resolution of 60000, an AGC target of 300% and a maximum injection time of 100 ms. For acquisition of MS2 spectra, the DIA isolation scheme consisted of 2 x 75 staggered windows (8 m/z width, 4 m/z offset) covering the 400–1000 m/z range. The resolution of MS2 was set to 15000, with an AGC target value of 1000%. The injection time was adjusted in automatic mode and the normalized collision energy was set at 30%.

Raw mass spectrometry files were processed with the DIA-NN software (version DIA-NN 1.9.2) for database search in library-free search mode. Data were searched against merged mouse entries of the UniProtKB protein database (Mouse reference proteome, 55,307 uniprot entries, downloaded on 2023-04-11). The carbamidomethylation of cysteines was set as a fixed modification, whereas oxidation of methionine and protein N-terminal acetylation were set as variable modifications. Specificity of trypsin digestion was set for cleavage after K or R, and one missed trypsin cleavage sites were allowed. Peptide length range was set to 7-30, precursor m/z range to 400-1000 and fragment ion range to 100-1700. Validation of the identifications was performed by setting the precursor false discovery rate (FDR) at 1%. Heuristic protein grouping was used, and protein groups were also validated at 1% FDR. Global normalization was enabled. The report.pg_matrix.tsv report was used to extract protein abundance metrics.

### Phylogenetic distribution and genomic context of CysK2

Homologs of the *M. tuberculosis* H37Rv proteins CysK2, CysK1, CysM, Cbs, and Cds1 were identified in the UniProt database by BLAST using the corresponding *M. tuberculosis* protein sequences as queries. Close CysK2 homologs were defined as proteins sharing >50% amino acid identity with *M. tuberculosis* CysK2 over the full-length sequence. UniProt accession numbers of all proteins used for phylogenetic analysis are provided in Table S3.

Phylogenetic analysis was performed with Phylogeny.fr in advanced mode (57), using the JTT substitution model. Branch support was assessed with the Approximate Likelihood-Ratio Test (aLRT), using the minimum of SH-like and Chi2-based support values. Trees were visualized and annotated using iTOL (58).

The genomic environment of *cysK2* was examined in the IMG/M database, and the presence of genes predicted to be involved in copper homeostasis in the vicinity of *cysK2*, including multicopper oxidases, copper chaperones, and copper-transporting P-type ATPases, was recorded and compared across species.

### Secondary ion mass spectrometry

For ^13^C labeling, *M. tuberculosis* was grown in 7H9 medium supplemented with ADC, 0.05% Tyloxapol, and 10 g/L ^13^C_3_-glycerol (Sigma; 99 atom % ^13^C). Bone marrow-derived macrophages were seeded on indium tin oxide (ITO)-coated glass coverslips (PGO, Präzisions Glas & Optik GmbH, Germany), which allowed monitoring of sample preparation and localization of infected cells by confocal microscopy prior to nanoSIMS analysis. Cells were infected with ^13^C-labeled bacteria at a multiplicity of infection of 1. After 4 h of incubation at 37°C in RPMI GlutaMAX HEPES (Gibco) supplemented with 10% FBS, infected cells were washed three times with DPBS and the medium was replaced with fresh medium containing 10% FBS and 200 µM CuCl_2_. At 3 days postinfection, cells were fixed in 4% paraformaldehyde for 2 h and washed three times with distilled water.

After drying in air at room temperature, the ITO glass slices were introduced into a NanoSIMS-50 Ion microprobe (CAMECA, Gennevilliers, France) operated in scanning mode (59, 60). The detailed analytical conditions for SIMS imaging were described previously (61). For this study, a tightly focused Cs^+^ primary ion beam steps over the surface of the sample and five secondary ion species (^12^C^-^, ^13^C^-^, ^12^C^14^N^-^, ^31^P^-^ and ^63^Cu^-^) were monitored in parallel to create images of these selected ion species. More specifically, the detection of ^63^Cu^-^, with a precise mass of 62.92959 amu, requires moderate mass resolution condition so as to prevent mass interference by neighbouring ion species (not only the very intense ^31^P^16^O_2_, 62,96358 amu, at a relative mass distance, M/ΔM, of 1852, but also an abundant ^31^P^32^S, 62,94583 amu, much closer, at a M/ΔM of 3875).

During the analysis, the primary beam intensity was 1 pA with a typical probe size of 100 nm and the raster size (field of view) was about 30 μm with an image definition of 512×512 pixels. The image acquisition was carried out using multiframe mode with a dwell time of 0.5 ms per pixel per frame, and up to 100 frames were acquired so as to allow the accumulation of the weak ^63^Cu^-^ signal. The overall counting time was then 50 ms per pixel and the total analysis time was about 3h40mn.

Image processing was performed using ImageJ software (62) by means of OpenMIMS plug-in (63). After raw data extraction, multiframe images were properly aligned using ^12^C^14^N^-^images as reference before a summed image was obtained for each ion species. A map of ^13^C atomic fraction was then derived from ^12^C^-^ and ^13^C^-^ images as ^13^C/(^12^C+^13^C). The location of bacteria inside the host cell was highlighted by high ^13^C composition. The distribution of ^31^P^-^ and ^63^Cu^-^ was shown based on the respective number of ion counts per pixel. These elemental and isotopic maps were displayed using Fire mode LUT (lookup table) in ImageJ. Further, color overlay images were generated using ImageJ to highlight the location of ^63^Cu^-^spots inside the bacteria, as well as the colocalization of ^63^Cu^-^ and ^31^P^-^ spots.

### RNA extraction and RT-qPCR

For bacterial RNA extraction, total RNA was isolated from 10 mL of *M. tuberculosis* cultures grown to logarithmic phase (OD_600_ = 0.8-1.0) using the RNeasy kit (Qiagen), following the manufacturer’s instructions with minor modifications. RNA samples were treated with DNase I (Thermo Scientific), and 500 ng of total RNA was reverse transcribed into cDNA using Moloney murine leukemia virus reverse transcriptase (M-MLV RT; Invitrogen) and random hexamer primers (Thermo Scientific). For eukaryotic RNA extraction, 1.5 × 10^6^ cells were resuspended in 1 mL TRIzol reagent. After addition of 200 µL chloroform and centrifugation, the aqueous phase was collected and total RNA was purified using the RNeasy Mini kit (Qiagen). cDNA was synthesized from 1 µg of total RNA using M-MLV reverse transcriptase (Invitrogen) and random hexamer oligonucleotides according to the manufacturer’s instructions.

Quantitative real-time PCR was performed using gene-specific primers (Table S2) and TB Green Premix Ex Taq (Takara) on an ABI Prism 7500 instrument (Applied Biosystems). Data were analyzed with 7500 software v2.0.6 (Applied Biosystems) and expressed as ΔCt values relative to the reference gene *rpoB* for bacterial samples or *Actb* for eukaryotic samples.

### RNA sequencing

The *M. tuberculosis* H37Rv strain was grown in triplicate in modified copper-free 7H9 medium supplemented with either no added copper, 20 µM CuSO_4_, or 100 µM CuSO_4_. Total bacterial RNA was extracted from 10 mL cultures harvested at logarithmic phase (OD_600_ = 0.8) using the RNeasy kit (Qiagen), as described above, and treated with DNase I (Thermo Scientific).

Bulk RNA sequencing was performed by Single Cell Discoveries (Utrecht, The Netherlands) using an adapted version of the VASA-seq protocol. Briefly, each total RNA sample was barcoded with CEL-seq primers during enzymatic lysis, followed by fragmentation, end repair, poly(A) tailing, reverse transcription, and second-strand synthesis. After second-strand synthesis, barcoded samples were pooled into a single library and amplified by *in vitro* transcription. Ribosomal amplified RNA was then depleted, after which adapter ligation, reverse transcription, and indexing PCR were performed to generate the final cDNA library. Libraries were sequenced in paired-end mode on an Illumina NovaSeq X Plus platform (read 1, 26 cycles; index read, 6 cycles; read 2, 60 cycles). Raw sequencing reads were processed with Cutadapt v4.9 to remove adapter sequences and homopolymers, then filtered with Ribodetector v0.3.1 to remove ribosomal reads. Processed reads were aligned to the *M. tuberculosis* H37Rv reference genome (assembly ASM19595v2) using STARsolo v2.7.11b with the parameter --soloFeatures GeneFull_Ex50pA.

Filtering of low-abundance reads, normalization, differential gene expression analysis, and z-score calculation were performed using edgeR (64). Differentially expressed genes were defined as those with an absolute log_2_(fold change) > 1 and a false discovery rate (FDR) < 0.05.

### Macrophages infection

Bone marrow cells were isolated from the femurs and tibias of C57BL/6 mice and differentiated into macrophages by culture for 7 days at 37°C in 5% CO_2_ in RPMI GlutaMAX HEPES medium (Gibco) supplemented with 10% FBS (Sigma), penicillin (10 U/mL), streptomycin (10 µg/mL), and 20 ng/mL M-CSF (PeproTech). After differentiation, macrophages were seeded in 24-well plates at 3 × 10^5^ cells per well on glass coverslips in antibiotic-free medium. Cells were infected with *M. tuberculosis* at a multiplicity of infection of 1 for 4 h at 37°C. Cells were then washed three times with DPBS, and fresh medium containing the indicated concentrations of CuSO_4_ was added. At 6 days postinfection, cells were washed twice with DPBS and fixed for 2 h at room temperature either in 4% paraformaldehyde in DPBS or, for Mrx1-roGFP reporter strains, in DPBS containing 4% paraformaldehyde and 10 mM N-ethylmaleimide. Fixed samples were washed three times with DPBS and analyzed by confocal laser scanning microscopy (Zeiss LSM710).

For Mrx1-roGFP strains, intracellular redox status was quantified as the ratio of oxidized to reduced probe forms, calculated on a pixel-by-pixel basis as the fluorescence intensity obtained with 405-nm excitation divided by that obtained with 488-nm excitation, with emission collected between 510 and 540 nm. Image processing and quantification were performed using ImageJ.

### Mouse infection for CFUs counting

All animal procedures were performed in facilities accredited under agreement F31555005 and by qualified personnel in accordance with French laws and regulations. Experimental protocols were approved by the French Ministry under authorization APAFIS #40827-2023020711364609 v8. Female C3HeB/FeJ or C57BL/6 mice, 6 to 8 weeks of age, were infected by aerosol using a DSI Buxco Inhalation Tower. Mice were exposed for 17 min to aerosols generated from a 5 mL bacterial suspension containing 2 × 10^6^ bacilli. At 7, 21, and 42 days postinfection, five mice per strain were euthanized and lungs were collected. Lung homogenates were plated on Middlebrook 7H11 agar supplemented with OADC, and bacterial burden was determined by CFU enumeration.

### Histology

C3HeB/FeJ mice were infected by aerosol with approximately 100 CFU of H37Rv, Δ*cysK2*, or the complemented strain Δ*cysK2*::*lpqS*-*cysK2*, each expressing the Mrx1-roGFP redox reporter. At 60 days postinfection, mice were euthanized by intraperitoneal injection of 50 µL Euthasol (400 mg/mL sodium pentobarbital). Lungs were perfused intratracheally with 1 mL of 2% agarose (UltraPure LMP Agarose; Invitrogen) and fixed for 24 h at 4°C in periodate-lysine-paraformaldehyde. Fixed lungs were cryoprotected by sequential incubation in 10%, 20%, and 30% sucrose prepared in 0.1 M phosphate buffer, embedded in optimal cutting temperature compound (OCT), and frozen. Sections of 20 µm thickness were cut on a cryostat and analyzed by confocal laser scanning microscopy (Zeiss LSM710).

### Disk diffusion assay

*M. tuberculosis* cultures were grown to exponential phase and diluted to an OD_600_ of 0.1 in 3 mL of prewarmed top agar (7H9 containing 6 mg/mL agar). The bacterial suspension was overlaid onto Middlebrook 7H10 agar plates. After solidification, a sterile filter paper disk was placed on the surface of the top agar and loaded with 5 µL of 1 M CuSO_4_. Plates were incubated under normoxic (21% O_2_) or hypoxic (10% O_2_) conditions, and inhibition zones were measured. Values were normalized to the inhibition zone obtained for the wild-type strain under normoxia.

### Metabolite Extraction

*M. tuberculosis* cultures were grown to an OD_600_ of 1.0 in 7H9 medium supplemented with 10% ADC and 0.05% Tyloxapol. Cultures were harvested by centrifugation and resuspended in DPBS at threefold the original concentration. For metabolite labelling experiments, 1 mL of each bacterial suspension was deposited onto 0.22-µm filters (mixed cellulose ester membrane; Millipore). Twenty-seven filters per strain were placed on minimal agar plates (KH_2_PO_4_, 0.5 g/L; MgSO_4_, 0.5 g/L; citric acid, 2 g/L; glycerol, 10 g/L; aspartate, 2 mM; agar, 1.5%; prepared in tap water) supplemented with 6 µM CuSO_4_ and 1 mM serine, with three filters placed on each plate, and incubated at 37°C. After 5 days, filters were transferred onto fresh minimal agar plates containing ^15^N-labeled serine and 6, 20, or 100 µM CuSO_4_ (nine filters per strain and per copper concentration). After 4 h of incubation, bacteria were recovered from the filters in 1 mL of quenching solution consisting of acetonitrile/methanol/water (40:40:20, vol/vol/vol). Cells were lysed with glass beads using a FastPrep-24 5G bead beater (MP Biomedicals) for three cycles of 45 s. Samples were centrifuged for 5 min, and the supernatant was collected and filtered twice through 0.2-µm Spin-X columns at 20,000 g for 15 min. Extracts were stored at −80°C until analysis. Protein concentration was determined using the Pierce BCA Protein Assay kit (Thermo Fisher Scientific).

### Liquid chromatography-mass spectrometry (LC-MS)

Metabolite extracts were analyzed using an Agilent 1290 Infinity II UHPLC system coupled to an Agilent 6545 LC/Q-TOF mass spectrometer. Chromatographic separation was performed on an Agilent InfinityLab Poroshell 120 HILIC-Z column (2.1 × 100 mm, 2.7 µm; p/n 675775-924), using a method optimized for polar acidic metabolites. The column temperature was maintained at 50°C.

A 10× stock solution of 100 mM ammonium acetate (pH 9.0) in water was prepared and used to generate the mobile phases. Mobile phase A consisted of 10 mM ammonium acetate in water (pH 9.0) supplemented with 5 µM Agilent InfinityLab deactivator additive (p/n 5191-4506). Mobile phase B consisted of 10 mM ammonium acetate (pH 9.0) in water/acetonitrile (10:90, vol/vol), also supplemented with 5 µM deactivator additive. The gradient was run at a flow rate of 0.25 mL/min as follows: 0-2 min, 96% B; 2-5.5 min, 88% B; 5.5-8.5 min, 88% B; 8.5-9 min, 86% B; 9-14 min, 86% B; 14-17 min, 82% B; 17-23 min, 65% B; 23-24 min, 65% B; 24-24.5 min, 96% B; 24.5-26 min, 96% B, followed by 3 min re-equilibration at 96% B.

Mass spectrometry was performed on an Agilent Accurate Mass 6545 QTOF instrument equipped with an electrospray ionization source operated in negative-ion mode. Dynamic mass-axis calibration was achieved by continuous infusion of a reference mass solution using an isocratic pump connected to the source. Instrument parameters were as follows: gas temperature, 225°C; drying gas flow, 13 L/min; sheath gas temperature, 350°C; sheath gas flow, 12 L/min; nebulizer pressure, 35 psi; capillary voltage, 3,500 V; nozzle voltage, 0 V; fragmentor voltage, 125 V; skimmer voltage, 45 V; and octupole 1 RF voltage, 750 V. Data were acquired in centroid 4 GHz extended dynamic range mode.

### Statistical analyses

Statistical analyses were performed using GraphPad Prism v10.2.3 (GraphPad Software). The statistical tests used for each experiment are indicated in the corresponding figure legends.

## Omics data availability

Proteomics and RNAseq data are being deposited in a public repository and will be accessible to readers.

## ACKNOWLEDGMENTS

We thank the Heran Darwin lab for comments on a draft version of this manuscript and sharing unpublished data about CysK2.

We gratefully acknowledge Marion Faucher, Bertille Voisin, and Pierre Dupuy for their valuable assistance with microbiology experiments, and Adeline Girel and Wladimir Malaga for their support in the construction of HoxB8 cell lines. We warmly thank Anne Gonzalez de Peredo for her careful verification of the proteomics data and their deposition in the PRIDE database. We acknowledge Sebastien Santini (CNRS/AMU IGS UMR 7256) and the PACA Bioinfo platform for the availability and management of the phylogeny.fr website used to reconstruct protein phylogeny. We thank the Wellcome Sanger Institute for providing us with a Cas9 nuclease-expressing mouse line that they generated using the hCas9 plasmid obtained from Addgene.

We acknowledge the Toulouse Réseau Imagerie imaging and cytometry facility, member of Genotoul and of the France-BioImaging infrastructure supported by the French National Research Agency (Grant ANR-24-INBS-0005 FBI BIOGEN), in particular Serge Mazeres and Eve Pitot for assistance with imaging, and Emmanuelle Näser for assistance with cytometry.

We also acknowledge the ANEXPLO facility for functional exploration, member of Genotoul and of the Celphedia infrastructure supported by the French National Research Agency (Grant ANR-10-INBS-04), in particular Malory Blasco, Flavie Moreau, and Céline Berrone for help with mouse care and infection experiments.

The Toulouse Proteomics facility is supported by the Région Occitanie, European funds (REACT-EU program), Toulouse Métropole, and by the French Ministry of Research with the “Infrastructures Nationales en Biologie et Santé” program to the Proteomics French Infrastructure, ProFI (ANR-10-INBS-08 & ANR-24-INBS-0015).

We acknowledge the CurieCoreTech, the network of technological facilities within Institut Curie, for the use of Curie-NanoSIMS. The Multimodal Imaging Center is a member of the French Infrastructures en Biologie Santé et Agronomie network.

This work was supported by CNRS, Université de Toulouse, the Ministry of Higher Education, Research and Space (PhD fellowship to W.L.M.), Agence Nationale de la Recherche sur le Syndrome d’immunodéficience acquise Maladies Infectieuses Émergentes (ANRS MIE, Grant ECTZ249116 to W.L.M.), Fondation pour la Recherche Médicale (Grant EQU202103012733 to O.N.), Fondation Bettencourt Schueller (Grant Explore-TB to O.N.), MSDAVENIR (Grant FIGHT TB to O.N.).

## AUTHOR CONTRIBUTIONS

W.L.M., F.L., T.-D.W., F. La., Y.P., and O.N. designed research; W.L.M., F.L., T.-D.W., Y.P., Y.R., E.V. and G.L.M. performed research; W.L.M., F.L., T.-D.W., Y.R., Y.P., and O.N. analyzed data; W.L.M., F.L., Y.P. and O.N. wrote the paper.

