## Supplementary Tables and Figures for "CysK2 couples copper sensing to metabolic and redox adaptation in *Mycobacterium tuberculosis*"

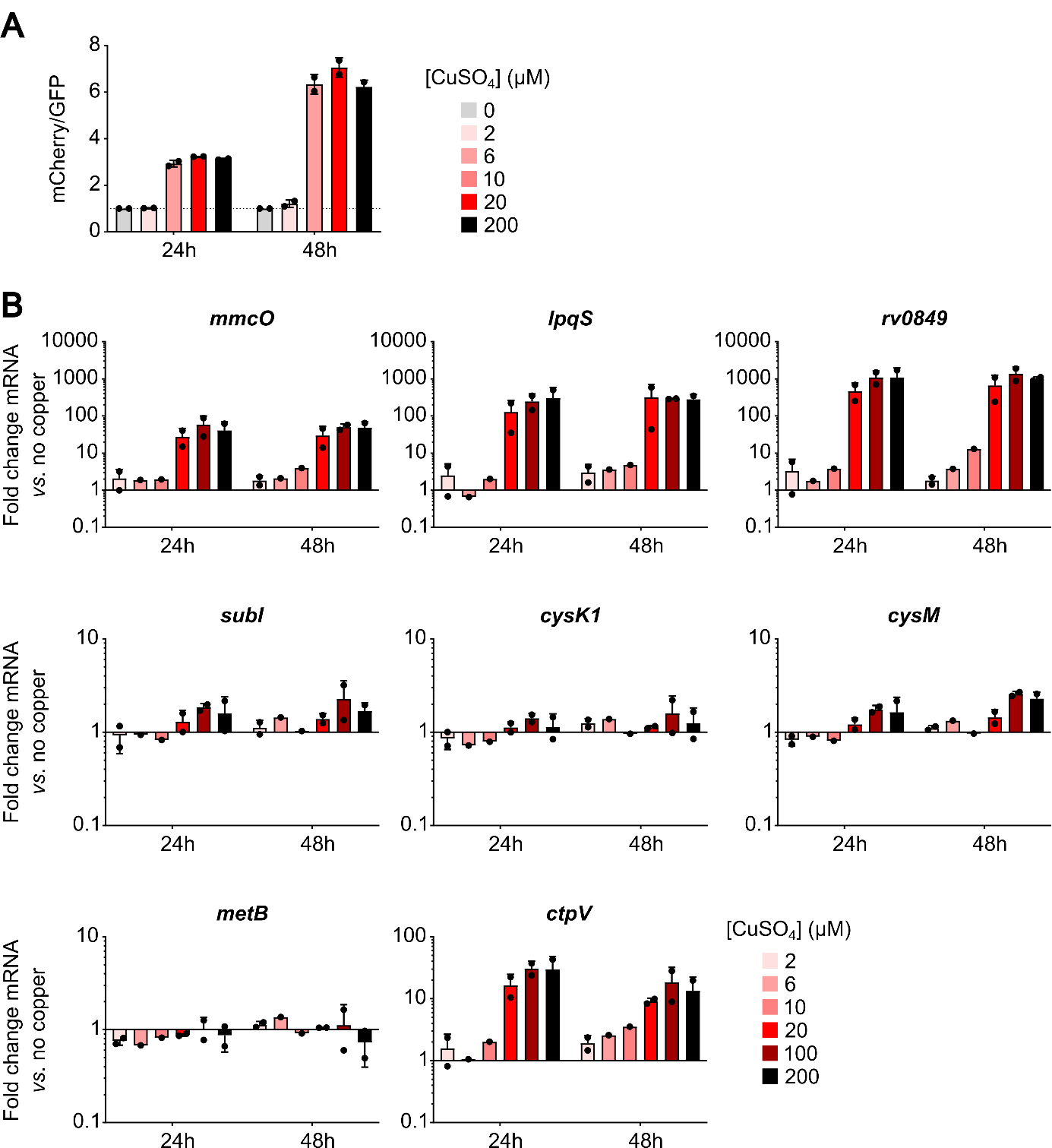

**Figure S1. Copper ions induce the expression of the *cysK2* locus but not of other genes related to sulfur metabolism.** *(A)* The P1-GFP *lpqSp*-mCherry reporter strain was incubated in PBS in the presence of 0, 2, 6, 20, or 200 µM CuSO₄ for 24 h or 48 h. The mCherry/GFP fluorescence ratio was then measured by flow cytometry *(B)* *M. tuberculosis* bacteria were incubated in M9 minimal medium in presence of 0, 2, 6, 10, 20, 100 or 200 µM CuSO_4_. After 24 or 48 hours, total RNAs were extracted and genes expression was quantified by RT-qPCR. Data represent the mean ± SD of fold change in genes expression compared to the reference gene *rpoB* and the condition 0 µM CuSO_4_ from two independent experiments.

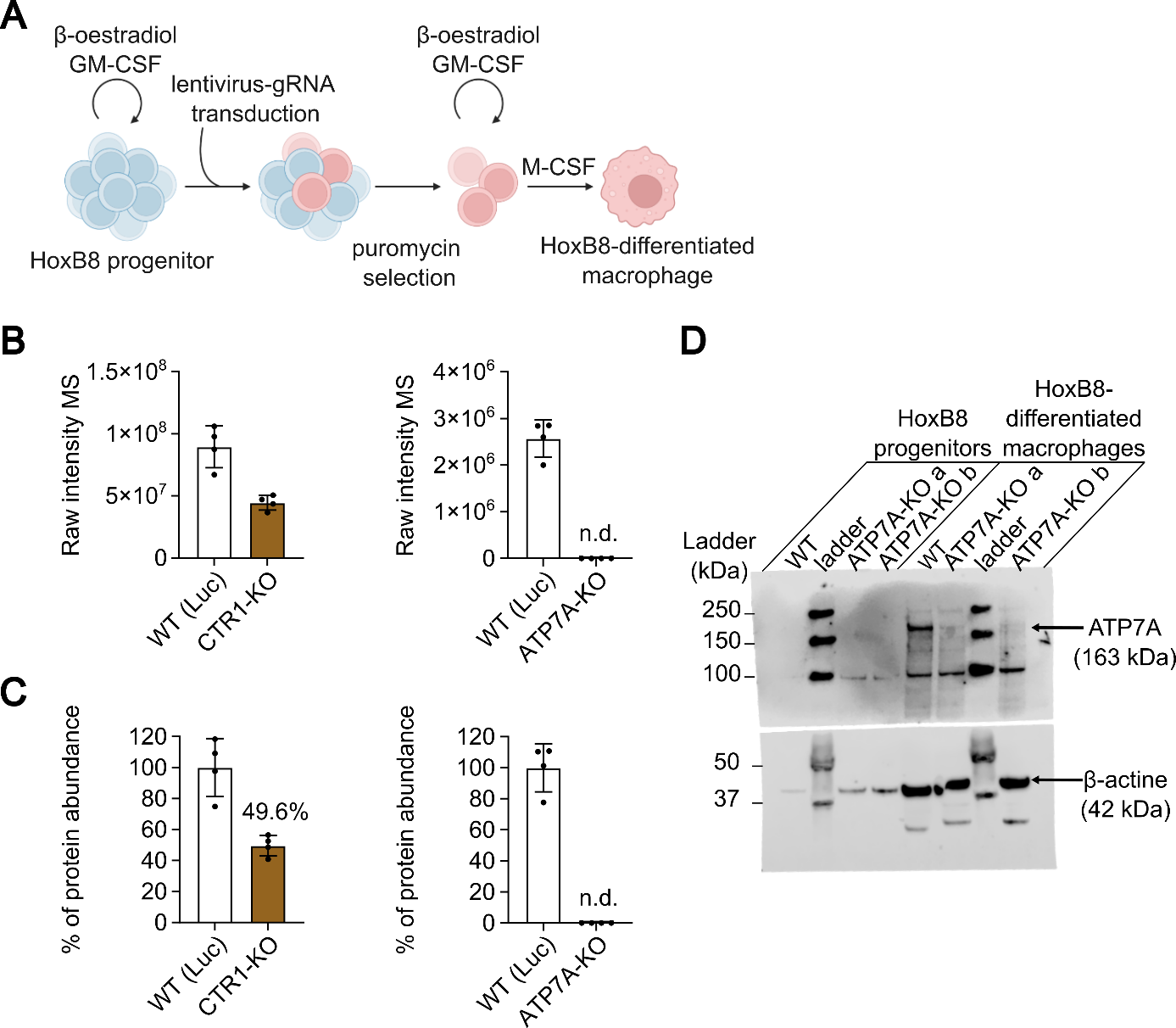

**Figure S2. Generation of CTR1- and ATP7A-deficient mouse macrophages**. *(A)* Schematic construction and differentiation of CRISPR-edited HoxB8 cell lines. Created in [https://BioRender.com](https://biorender.com/). *(B) (C)* Quantification of CTR1 and ATP7A proteins abundance in WT (Luc) and corresponding CRISPR-edited HoxB8-differentiated macrophages by proteomics. Data represent raw intensity *(B)* and percentage expression relative to WT expression *(C)*. *(D)* Western blot analysis of ATP7A in WT and ATP7A-KO clone a (used in this study) and clone b HoxB8 progenitors or HoxB8-differentiated macrophages.

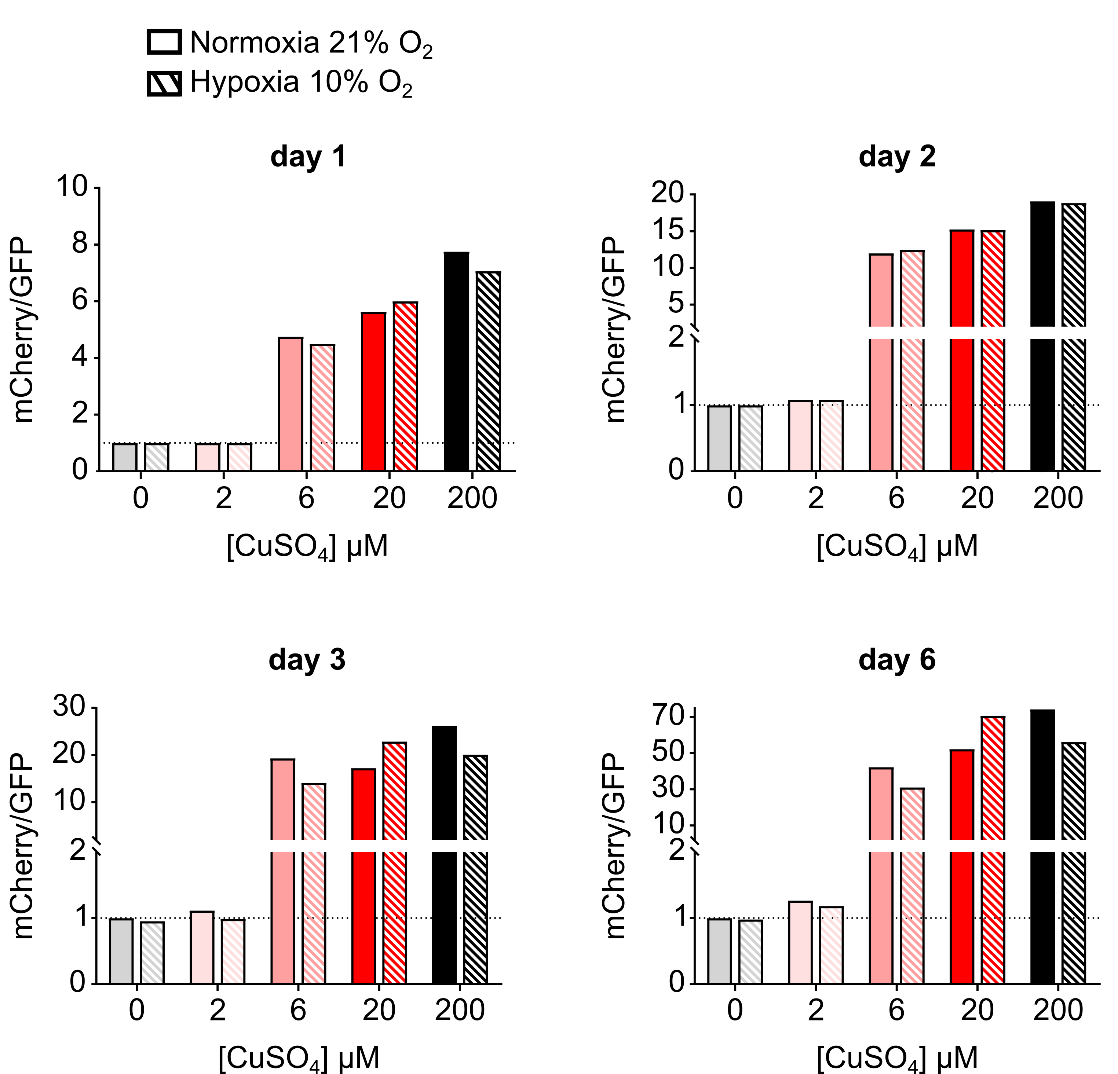

**Figure S3. Hypoxia does not boost copper-mediated *cysK2* induction in *M. tuberculosis in vitro.*** *M. tuberculosis* P1-GFP *lpqSp*-mCherry reporter strain was incubated in PBS in presence of 0, 2, 6, 20 or 200 µM CuSO_4_. At different time post-incubation, bacteria were fixed and mCherry/GFP ratio was quantified by flow cytometry and normalized to the condition 0 µM CuSO_4_. Data come from two independent experiments (experiment 1: days 1 and 2; experiment 2: days 3 and 6).

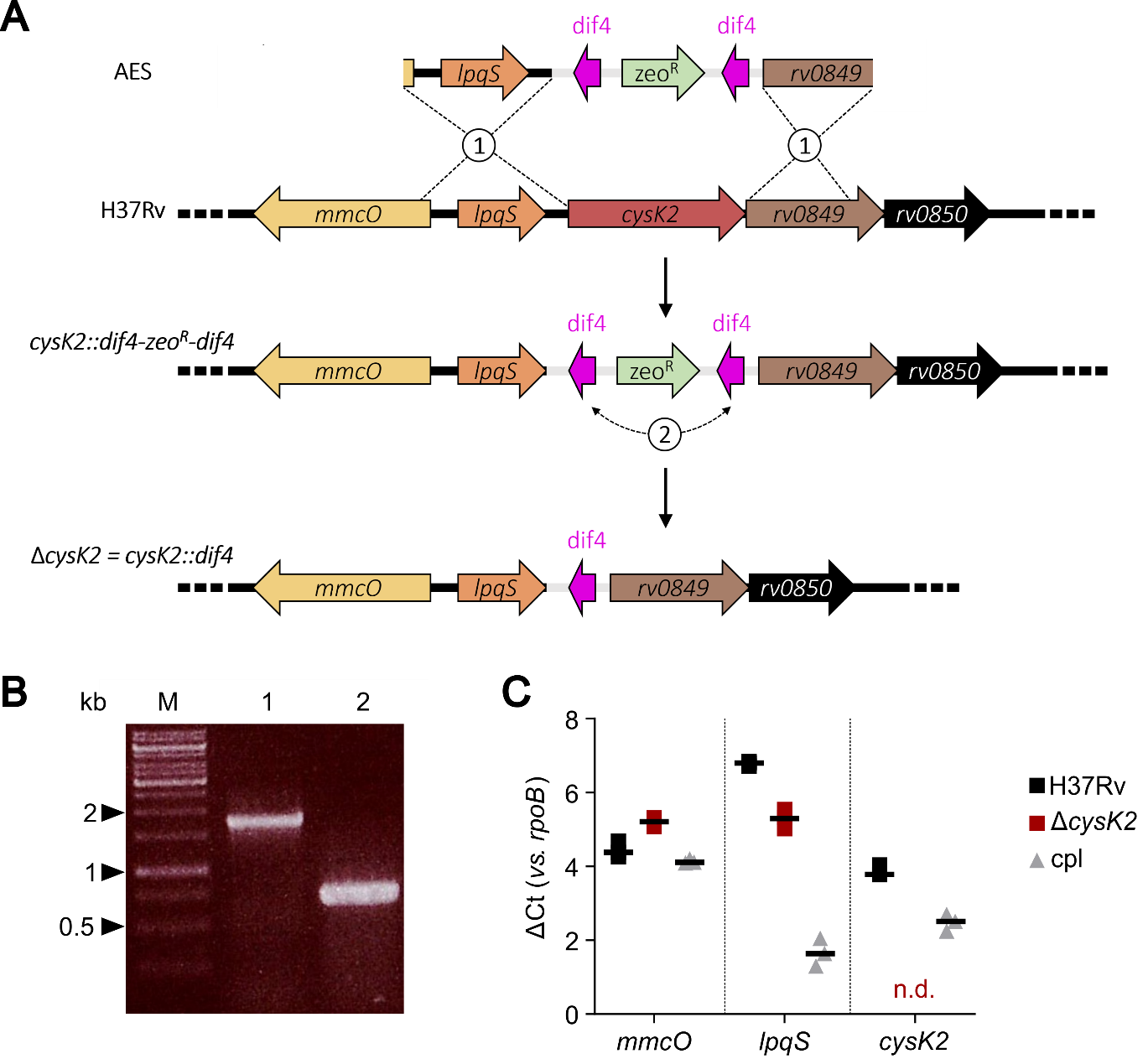

**Fig. S4.** **Construction and validation of *M. tuberculosis* ∆*cysK2* deleted strain and complemented strain**. *(A)* Schematic representation of ∆*cysK2* mutant construction strategy. The allelic exchange substrate (AES) consisting in a zeocin resistance cassette flanked by two identical dif sites and the up- and downstream fragments of the target *cysK2* gene recombines with H37Rv genome and replaces the *cysK2* gene resulting in a ∆*cysK2::dif4-zeo-dif4* mutant (1). The two dif sites recombine, leading to the loss of the zeocin cassette and to the ∆*cysK2::dif4*=∆*cysK2* mutant (2). *(B)* Confirmation of *cysK2* deletion. PCR was done with primers amplifying a 1.7 kb fragment in the wild-type strain (lane 1) and a 0.7 kb fragment in the ∆*cysK2* mutant (lane 2). M: molecular size marker. *(C)* RT-qPCR quantification (∆Ct values *vs. rpoB* reference gene) of *mmcO*, *lpqS* and *cysK2* expression in *M. tuberculosis* H37Rv, ∆*cysK2* mutant and ∆*cysK2::lpqS-cysK2* complemented strain. No *cysK2* complementary DNA was amplified from ∆*cysK2* extract. Data represent mean of three technical replicates.

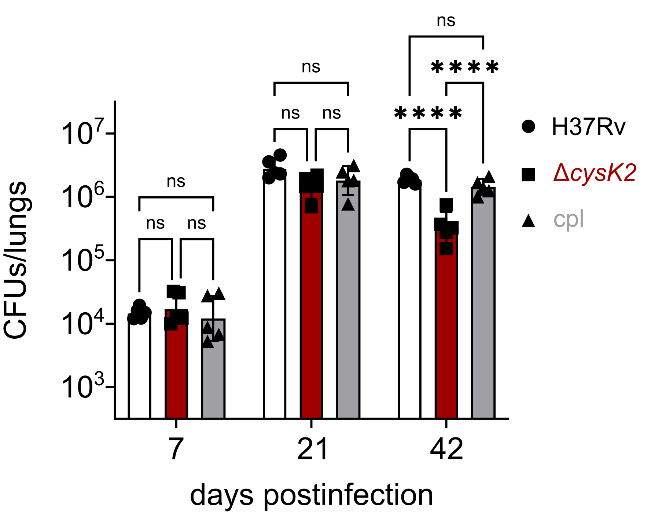

**Figure S5. CysK2 contributes to *M. tuberculosis* survival in C57BL/6**. Mice were infected via aerosol with H37Rv, ∆*cysK2* or complemented strains (100 CFUs per mice). Lungs were collected at 7, 21, and 42 days post-infection, and bacterial burden was quantified by CFU counting. Data represent mean ± SD for n=5 mice per strain. Statistical analysis: two-way ANOVA with Tukey’s post-hoc test on log-transformed data.

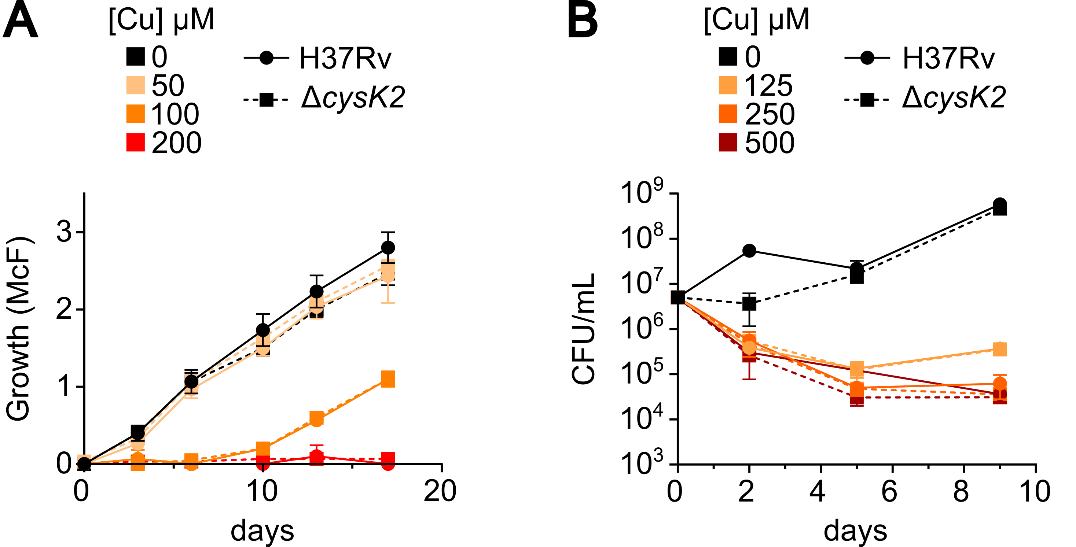

**Figure S6. The Δ*cysK2* mutant is not hypersensitive to copper in liquid medium as compared to the wild-type strain.** Both strains were cultured in 7H9 liquid medium in the presence of copper. *(A)* Bacterial growth was measured over a 17-day period (McFarland). *(B)* Survival of bacteria (CFUs count on 7H11 medium) in the presence of copper over a 9-day period.

**Table S1: Distribution of CysK2 orthologues among orders of Actinomycetes.**

| **Order (Class Actinomycetes)** | **Number of CysK2 homologues in UNIPROT** |
| --- | --- |
| Acidothermales | ND |
| Actinomycetales | ND |
| Bifidobacteriales | ND |
| Catenulisporales | 1 |
| Cryptosporangiales | ND |
| Frankiales | 3 |
| Geodermatophilales | 0 |
| Glycomycetales | 4 |
| Jatrophihabitantales | ND |
| Jiangellales | 1 |
| Kineosporiales | ND |
| Kitasatosporales | >100 |
| Micrococcales | 2 |
| Micromonosporales | 32 |
| Motilibacterales | ND |
| Mycobacteriales | >100 |
| Nakamurellales | ND |
| Propionibacteriales | 4 |
| Pseudonocardiales | >100 |
| Sporichthyales | ND |
| Streptosporangiales | >100 |

(*ND, not detected)

**Table S2: Primers used in this study**

| **Oligonucleotide** | **Sequence 5’->3’** | **Purpose** |
| --- | --- | --- |
| *Rv0848_up-Fw* | *gcgggatatcactagtgcgcgtgctgacgatcggcc* | ΔcysK2 mutant |
| *Rv0848_up-Rv* | *cagtcgatccacgtggagcaaggtgcggtgcgcccg* |  |
| *Rv0848_down-Fw* | *ccactgagcgtcagacccacgtgctcaatgggagcgcgcgctat* |  |
| *Rv0848_down-Rv* | *gcgttgggagctctcccagcatttcgaaaggaaaca* |  |
| *lpqS_up-Fw_IF* | *gccacgtggaaagcgggcacggggtcagcgccaggcggtg* | ΔcysK2 Cpl strain |
| *lpqS_up-Rv_IF* | *caccgcctggcgctgaccccgtgcccgctttccacgtggc* |  |
| *attB2-SD-rvcysK2* | *ggggacagctttcttgtacaaagtggaggaagacaggctgcccatgaggtcgcggcagacccg* |  |
| *attB3-rvcysK2_Rv* | *ggggacaactttgtataataaagttgttacgacaccacctgggttg* |  |
| *prom-rv0847-rv0850Fw* | *ttgtacaaaaaagcaggcttcggcgtcgtagtggcctgg* | *pGMCS-P1-GFP-P_rv0847-rv0850_-mCherry vector* |
| *IF-prom-rv0847-rv0850Rv* | *ctttgtacaagaaagctgggtcacgtgcccgctttccacg* |  |
|  | **RT-qPCR primers** |  |
| *rpoB-Fw* | *TCGTTCTCTGACCCTCGTTTC* |  |
| *rpoB-Rv* | *ACGTGCCCTTCTCGGTCATCA* |  |
| *cysK2-Fw* | *CGAGTATTTATCCGCGCAAT* |  |
| *cysK2-Rv* | *GTGTCGAAGTAGCGTTGTGG* |  |
| *Atp7a-Fw* | *AATTGCCATAGGCACAGGCAC* |  |
| *Atp7a-Rv* | *ATGGGCAGGAAAACTCCAGCA* |  |
| *Ctr1-Fw* | *ACATTACCATGCCACCTCACCA* |  |
| *Ctr1-Rv* | *CGAAGGCTCCAGCCATTTCTCCA* |  |
| *ActinB-Fw* | *CTAAGGCCAACCGTGAAAAG* |  |
| *ActinB-Rv* | *ACCAGAGGCATACAGGGACA* |  |

**Table S3: Accession numbers of proteins used for the phylogenetic analysis shown in Fig. 3A.**

| **Species** | **UniProt Accession Number** |
| --- | --- |
| *M. smegmatis* ATCC 700084 | A0R1X2_MYCS2 (CysM); A0R5I5_MYCS2 (Cds1); A0R2X8_MYCS2 (Cbs) |
| *M. abscessus* ATCC 19977 | B1MGL6_MYCA9 (CysK2); B1MFV5_MYCA9 (Cds1); B1MKR9_MYCA9 (CysK2); B1MLY9_MYCA9 (Cbs); B1MKJ1_MYCA9 |
| *M. marinum* ATCC BAA-535 | B2HL85_MYCMM (CysK1); B2HF70_MYCMM (CysM); B2HG60_MYCMM (CysK2); B2HK82_MYCMM (CysK1); B2HT76_MYCMM (Cbs) |
| *M. kansasii* ATCC 12478 | A0A7G1IC04_MYCKA; A0A1V3X6J4_MYCKA (CysK2); A0A7G1I8R6_MYCKA; A0A1V3XHI2_MYCKA (Cbs); A0A7G1IAD3_MYCKA |
| *M. fortuitum* ATCC 6841 | A0A0N9XJC1_MYCFO; A0A0N9XGY7_MYCFO (CysM); A0A0N9YME9_MYCFO (Cds1) |
| *M. vaccae* ATCC 25954 | K0VGZ0_MYCVA (Cds1); K0UXW7_MYCVA; K0UZ64_MYCVA (Cbs) |
| *M. gordonae* ATCC 14470 | A0A1X1VMA1_MYCGO (Cds1); A0A1A6B6N8_MYCGO; A0A1A6B7I0_MYCGO (Cbs) |
| *M. avium* ATCC BAA-968 | Q73Y35_MYCPA (CysK1); Q744F1_MYCPA (Cds1); Q741R3_MYCPA (CysM) |
| *M. terrae* ATCC 15755 | A0AAD1HYC3_9MYCO; A0AAD1HXV6_9MYCO (Cbs) |
| *M. aurum* ATCC 23366 | A0A3S4RRU7_MYCAU (CysM); A0A448IYF0_MYCAU (Cbs) |
